# A 2-Hydroxybutyrate- mediated feedback loop regulates muscular fatigue

**DOI:** 10.1101/2023.10.16.562554

**Authors:** Brennan J Wadsworth, Marina Leiwe, Eleanor A Minogue, Pedro P Cunha, Viktor Engman, Carolin Brombach, Christos Asvestis, Shiv K Sah-Teli, Emilia Marklund, Peppi Koivunen, Jorge L Ruas, Helene Rundqvist, Johanna T Lanner, Randall S Johnson

**Author notes:** **Corresponding Author:** Dr. Randall S Johnson.

## Abstract

The metabolite 2-hydroxybutyrate (2HB) is produced by skeletal muscle acutely during exercise and persists for several hours in the blood post-exertion. We show here that 2HB directly inhibits branched-chain aminotransferase enzymes, and that this inhibition in turn triggers a SIRT4-dependent shift in the compartmental abundance of protein ADP-ribosylation. The 2HB-induced decrease in nuclear protein ADP-ribosylation leads to a C/EBPβ mediated transcriptional response in the branched-chain amino acid degradation pathway. This response to 2HB exposure leads to an improved oxidative capacity both *in vitro* and *in vivo*, with the latter mimicking the effects of exercise training on whole body metabolism. Thus, we show here that 2-HB production by skeletal muscle represents a novel mechanism for the modification of metabolism by exercise.

## Introduction

Physical exertion is a metabolic challenge that requires physiological responses throughout the body, including a dramatic change to metabolic flux patterns^1^. During exercise, metabolic end products accumulate and are released into the circulation; prototypical examples of such molecules include hypoxanthine and lactate. Metabolic intermediates of pathways with increased flux during exercise may also exhibit increased levels in the circulation post-exercise, particularly as an indication of pathway saturation^2^. Accumulation of individual exercise-induced metabolites may lead to a diverse range of cellular responses in different tissues, with acute and lasting effects on physiology. Such mechanisms have been defined for succinate^3^, lactate^4^, hypoxanthine^5^, and other exerkines^1^, demonstrating that the exercise-induced metabolome includes factors with significant physiological effects.

Exercise-induced lactate and succinate exert effects on physiology despite a relatively rapid return to baseline levels during recovery post-exercise. Conversely, 2-hydroxybutyrate (2HB, also known as α-hydroxybutyrate) is an exercise-induced metabolite with plasma concentrations that continue to increase for at least 3 hours after exertion^6–9^. Production of the majority of 2HB in circulation has been attributed to the liver, although 2HB accumulates in many other tissues post-exercise^2^. 2HB is also present in disease states, as untargeted metabolomics analyses have identified 2HB as a metabolic marker for infection severity^10^, and as an indicator of metabolic disorders^11–13^. Cellular responses to accumulated 2HB have not been characterized to date, although Sato, et al., report that exogenous administration of 2HB caused an acute shift in mouse whole body metabolism, including a reduction in the respiratory exchange ratio (RER), and increased blood glucose^2^; indicating that 2HB has the potential to alter metabolism.

2HB is the product of a lactate dehydrogenase (LDH)-mediated reduction of 2-ketobutyrate (2KB), which itself is a byproduct of endogenous cysteine synthesis via the transsulfuration pathway. Oxidation of 2KB is not well characterized beyond isolated enzyme assays; these show it to be a substrate for the branched chain keto acid dehydrogenase complex (BCKDH)^14^. BCKDH converts 2KB to propionyl-CoA, which can be re-carboxylated by propionyl-CoA carboxylase (PCC), producing the tri-carboxylic acid (TCA) cycle intermediate succinyl-CoA^15^. Mammalian cells are not capable of reverse transsulfuration, i.e., production of methionine from cysteine; thus, the metabolic fate of 2KB is driven by the balance between mitochondrial oxidation or reduction to 2HB. Importantly, the metabolism of exogenous 2HB has not been described. Based upon the current literature, the fate of 2HB depends on the capacity for LDH-dependent oxidation of 2HB, and thus, 2KB metabolic homeostasis.

2HB is an alpha-hydroxy carboxylic acid similar to 2-hydroxyglutarate (2HG). The latter is a dicarboxylic acid, and an established physiological competitor of α-ketoglutarate (αKG). Both the *S*- and *R*- enantiomers of 2HG are known to competitively inhibit αKG-dependent enzymatic reactions, including the branched chain amino-transferase (BCAT) enzymes, and the much broader family of αKG-dependent dioxygenases (αKGDD)^16^. αKGDDs include regulators of the cellular responses to hypoxia, the prolyl hydroxylase (PHD) and factor inhibiting HIF (FIH), and epigenetic regulators such as lysine demethylases (KDM), among others. Regulation of these targets by endogenous or exogenous *S*-2HG is sufficient to alter numerous cell fate decisions, e.g., the differentiation of cytotoxic CD8 T cells^17,18^. *R*-2HG inhibition of BCAT is reported to reduce metabolism of branched chain amino acids (BCAA), leading to reduced levels of glutamate and glutathione^19^. As an alpha-hydroxy acid, we hypothesised that 2HB may be a novel competitive inhibitor of αKGDDs or BCATs.

In this work we investigate the physiological and molecular role of 2HB. We find that cells treated with 2HB demonstrate a metabolic response that leads to transcriptional regulation of BCAA degradation enzymes via a BCAT2 and SIRT4-dependent shift in compartmental protein ADP-ribosylation (ADPr). The reduced ADPr in the nuclear compartment following 2HB treatment promotes binding of the transcription factor CCAAT enhancer binding protein beta (C/EBPβ) to the promoters of BCAA degradation genes. We find that repeated administration of 2HB to mice increases oxidative capacity in exercise tests that replicates the effect of exercise training. Finally, we find repeated 2HB treatment gives rise to an improved resistance to fatigue in oxidative skeletal muscle, consistent with the improvement to oxidative metabolism. Overall, we find that 2HB induces a range of cellular responses that are consistent with it being a feedback signal from exertion, one that leads to increased capacity in the BCAA degradation pathway and improved exercise performance.

## Results

### Daily 2HB treatment accelerates the metabolic response to exercise in mice

As described above, physical exertion induces a persistent increase to circulating 2HB. To investigate the role of 2HB in whole animal metabolism, we first subjected mice to daily incremental exhaustive exercise tests for five consecutive days. Immediately following each test, mice were injected with 1 mmol/kg of Na2HB, or an equivalent dose of NaCl as a control (Fig.1A). Mouse basal VO_2_, maximum ΔVO_2_ (change in VO_2_ relative to basal VO_2_), RER, and ΔVO_2_ throughout exercise were similar across treatment groups on the first day, prior to group randomization (Fig.1B-D,SFig.1A). Median mouse ΔVO_2_max was 507 mL/kg/hr on the first day (Fig.1C) and mice tended to reach exhaustion during the 21 m/min phase (Fig.1E). Beyond 18 m/min, mice either further increased their VO_2_ or reached exhaustion. This is demonstrated by the lack of correlation between ΔVO_2_ during 18 m/min and time-to-exhaustion, but a significant correlation between either ΔVO_2_ during 21 m/min or maximum ΔVO_2_ and time-to-exhaustion (SFig.1B). These data demonstrate that increased oxidative capacity predicts improved performance in incremental exercise tests.

**Figure 1.**
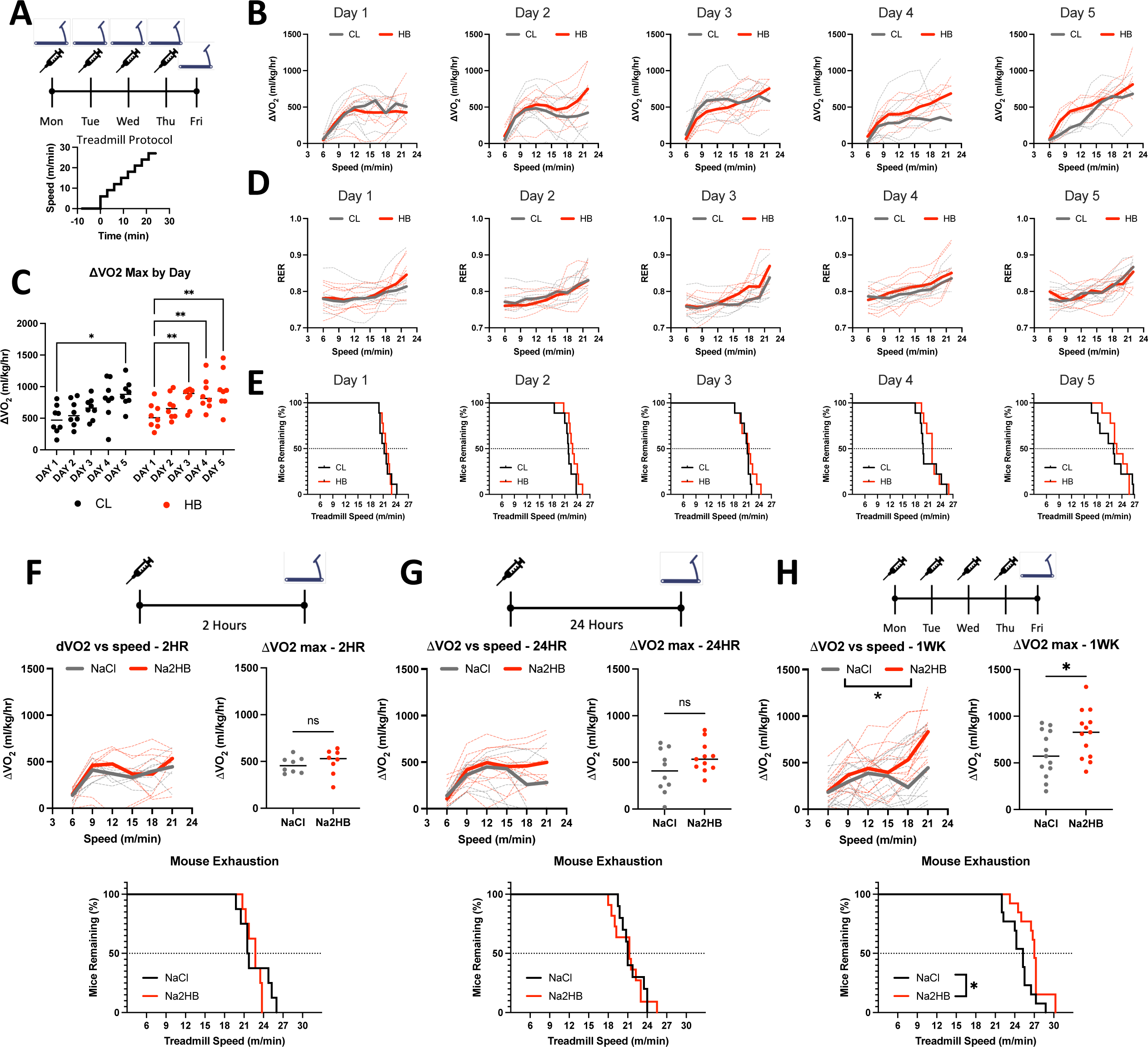
(A) Mice were subjected to five days of daily incremental exercise tests to exhaustion followed immediately by injection with 1 mmol/kg of NaCl or Na2HB, N = 8. (B) Change in VO_2_ relative to baseline (ΔVO_2_) versus treadmill speed. (C) Maximum ΔVO_2_ compared with Day 1. Dunnett’s multiple comparisons test, * p < 0.05, ** p < 0.01. (D) Respiratory exchange ratio (RER) versus treadmill speed. (E) Time-to-exhaustion. Mice were subject to a single incremental exercise test (F) 2 hours (N = 8) or (G) 24 hours (N = 11) after a single dose, or (H) after four daily doses (N = 13) of NaCl or Na2HB. For B,D, and F-H, solid lines indicate median values and dashed lines show each replicate. Median plots clipped at 21 m/min due to mouse dropout, although mice may continue up to 30 m/min. Speed vs ΔVO_2_ plot shows two-way ANOVA main effect * p < 0.05, ΔVO_2_ max comparison shows student’s t-test * p < 0.05. Time-to-exhaustion plots show Log-Rank test, * p < 0.05.

Mouse ΔVO_2_max tended to increase with subsequent exercise tests and had significantly improved to a median of 877 mL/kg/hr in the control group by day 5 (Fig.1C). Accordingly, progressively more mice reached the 24 m/min phase before exhaustion (Fig.1E). Mice treated with 2HB showed improvement to ΔVO_2_max in subsequent exercise tests as early as day 3, and increased ΔVO_2_max up to 930 mL/kg/hr on day 5 (Fig.1C). These data demonstrate a benefit to exercise performance from a brief one-week protocol of exercise training in mice and suggest that combination with exogenous 2HB may accelerate this effect.

### Daily 2HB treatment recapitulates the benefits of exercise training on oxidative capacity

We next investigated the effects of exogenous 2HB alone. Firstly, we assessed the effects of acute administration of 1 mmol/kg of Na2HB. For mice at rest, we observed no effect of exogenous 2HB on oxygen consumption or RER compared with NaCl control over the first 60 minutes, consistent with previous reports for this dose (SFig.1C,D)^2^. We next administered 2HB 2 hours prior to an exhaustive exercise test. Both 2HB-treated and control mice reached a ΔVO_2_max of approximately 500 mL/kg/hr, with no difference in RER, time-to-exhaustion, resting VO_2_, or ΔVO_2_ during the exercise tests (Fig.1F, SFig.1E,F). These data indicate that exogenous 2HB at a dose of 1 mmol/kg does not acutely alter resting metabolism or oxygen consumption during exercise.

Next, we investigated whether treatment with 2HB would alter exercise performance on subsequent days. First, a cohort of mice was treated with 2HB 24 hours prior to an exhaustive exercise test. In these mice there was again no difference between groups in terms of resting VO_2_, ΔVO_2_ or RER during the exercise test, time-to-exhaustion, or ΔVO_2_max (Fig.1G, SFig.1E,F). When mice were treated with 2HB each day for four days with the exhaustive exercise test on the fifth day, 2HB- treated mice exhibited an increased ΔVO2 during the exercise test, with differences more apparent at greater treadmill speeds (Fig 1H). Oxygen consumption at higher treadmill speeds correlated with greater time-to-exhaustion, indicating effective oxidative capacity as opposed to metabolic inefficiency (SFig.1G). Mice treated with 2HB exhibited a greater ΔVO_2_max and greater time-to-exhaustion, but had similar basal VO_2_ and RER levels throughout exercise, relative to exercising NaCl-treated controls (Fig.1H, SFig.1E,F). Control mice exhibited a median ΔVO_2_max of 572 ml/kg/hr. However, 2HB-treated mice exhibited a median ΔVO_2_max of 828 mL/kg/hr, similar to exercise-trained mice (Fig.1C,H). These data indicate that repeated 2HB treatment replicates the improvement to mouse oxidative capacity and exercise performance produced from daily exercise training.

### 2HB levels are correlated with metabolic indicators of BCAA degradation in murine datasets

To determine which metabolic pathways 2HB may affect, we were next interested in determining which metabolites are most correlated with 2HB post-exercise. We analyzed two published datasets reporting untargeted metabolomics in mice subjected to a treadmill-based exercise protocol, or sham control^2,8^. Both studies observe a strong increase to 2HB in the circulation and in all assessed organs post-exercise (SFig.2A). A correlation analysis demonstrates a reliable positive correlation between circulating 2HB and markers of BCAA metabolism (Fig.2A). Overrepresentation analysis of BCAA-related metabolites positively correlated with 2HB post-exercise yielded p values of 3.8×10^-5^ and 1.6×10^-2^ for the two studies. These data suggest that accumulation of 2HB is correlated to the metabolic flux through the BCAA degradation pathway.

**Figure 2.**
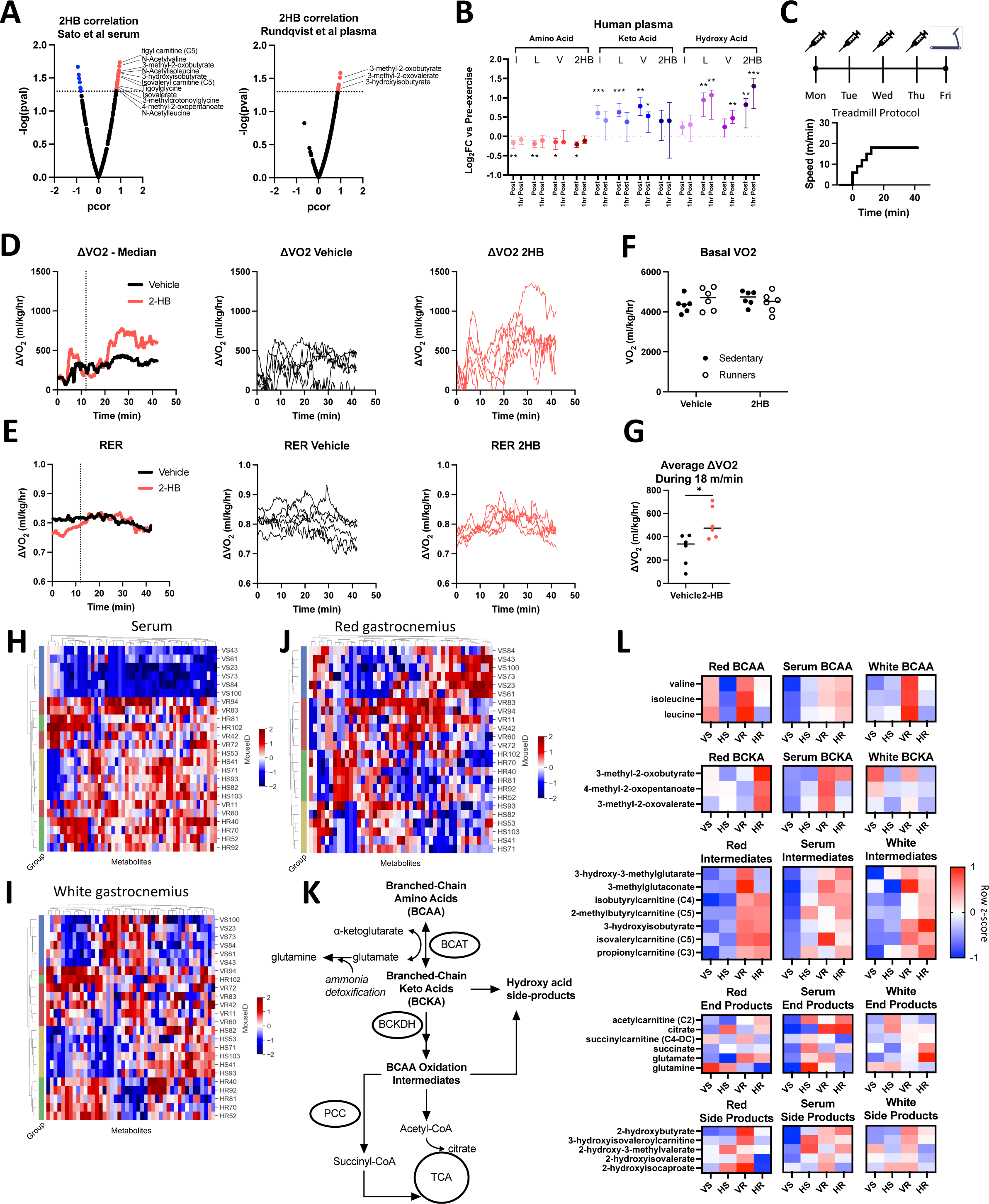
(A) Volcano plots display partial correlation coefficients for metabolites present in the blood that correlate with 2HB levels in mice that have undergone exercise. BCAA degradation metabolites indicated. Data in A are from Sato et al. 2022 (*left*) and Rundqvist et al. 2021 (*right*). (B) Levels of amino acid, keto-acid, and hydroxy acid versions of the BCAA and 2HB are displayed for human subjects comparing pre-exercise measures with those immediately post exercise and one-hour post-exercise. Data from Rundqvist et al. 2021, N = 8, median ± interquartile range. Dunnett’s multiple comparisons test made against pre-exercise levels, *** p< 0.001, ** p< 0.01, * p< 0.05. (C) Mice were treated with 1 mmol/kg of NaCl or Na2HB for four days prior to a single endurance exercise protocol consisting of a gradual warm-up and 30 minutes of running at 18 m/min. Sham control mice placed on motionless treadmill. (D) ΔVO_2_ and (E) RER plots, Left-to-Right: group medians, vehicle-treated, and 2HB-treated. (F) No effect of 2HB on basal VO_2_. (G) 2HB-treated mice show increased average ΔVO_2_. Student’s t-test, * p < 0.05. Untargeted metabolomics was conducted on serum, red gastrocnemius, and white gastrocnemius from mice in figures C-G. Heat maps of top 50 most differentially abundant metabolites in (H) serum, (I) white gastrocnemius, and (J) red gastrocnemius. Unique mouse ID displayed with group abbreviations: ‘VS’ vehicle-sedentary, ‘HS’ 2HB-sedentary, ‘VR’ vehicle-runner, ‘HR’ 2HB-runner. (K) BCAA degradation pathway. (L) Heat map displays metabolites in the BCAA degradation pathway. For D-L, N = 6. Heat maps display standardized row medians.

### Human exercise studies demonstrate the persistence of serum 2HB post-exercise

We further investigated this pattern by assessing the plasma metabolomics of human subjects in a previously published dataset from our group. In this study, blood was sampled prior to exercise, immediately post-exercise, and then one-hour post-exercise. Few markers of BCAA degradation are available in the human dataset, so we focused on the amino acid, the keto acid, and the hydroxy acid versions of leucine, isoleucine, valine, and 2HB. Transamination of BCAA to the respective keto acid is the first step towards mitochondrial oxidation. In line with this, BCAA levels are reduced immediately post-exercise compared with pre-exercise levels, while keto acids and hydroxy acids are increased (Fig.2B). The accumulation of 2HB alongside increased BCAA degradation is consistent with a model wherein BCAAs compete with 2KB for early oxidation steps, e.g., the BCKDH and PCC-mediated reactions.

Indicating a return to homeostasis, one-hour after completion of exercise, BCAAs increase and keto acids decrease towards pre-exercise levels (Fig.2B). The hydroxy acids are intriguing, as these remain significantly greater than pre-exercise levels even one-hour after completion of exercise, suggesting slow clearance of all members of this metabolite group, including 2HB. Among the hydroxy acids, 2HB is the standout, with the greatest fold-increase at the one-hour time point and the greatest absolute abundance in both human and mouse datasets^8^. Collectively, these data describe a positive correlation for the production of 2HB in proportion with production of branched chain keto acids (BCKA) during exercise. The slow clearance of 2HB and the branched chain hydroxy acids (BCHA) represent a possible indication of saturation in the capacity for the oxidation of 2KB and BCKAs.

### Repeated 2HB treatment alters BCAA metabolite balance following exercise

We next investigated whether repeated treatment of 2HB for four days would alter BCAA metabolism during an exercise test on the fifth day. To limit mouse dropout from exhaustion, the exercise protocol increased speed only up to 18m/min. This speed was held for 30 minutes for all mice (Fig.2C). There was no effect of 2HB treatment on basal VO_2_, or on RER throughout the exercise protocol (Fig.2D-F). 2HB-treated mice tended to spend more time running on the treadmill, indicated by an increased ΔVO_2_ during the 18m/min segment of the exercise protocol (Fig.2D,G). These data demonstrate again that repeated 2HB treatment increases oxidative capacity during exercise.

We hypothesized that the change to exercise performance, in both the incremental exercise tests to exhaustion and moderate exercise tests, was in part due to a change in skeletal muscle metabolism during exercise. Untargeted metabolomics of mouse serum, red gastrocnemius, and white gastrocnemius collected immediately post-exercise shows modest clustering of mice based on treatment group in the serum, separating vehicle-treated sedentary controls from other treatment groups (Fig.2H). Conversely, we observe strong clustering in the red and white gastrocnemius, delineating each experimental group (Fig.2I,J). These data begin to suggest that skeletal muscle is a site of metabolic regulation by 2HB.

When we visualize the data trends in the BCAA metabolic pathway within each tissue (Fig.2K,L), a shift in the BCAA, BCKA, and BCHA appears to be exercise and 2HB dependent; specifically 2HB treatment appears to reduce BCAA levels in red gastrocnemius at rest and during exercise, but increase BCKA and decrease the hydroxy acids during exercise (Fig.2L). The levels of intermediates within the BCAA degradation pathway increase in each compartment with exercise, with no strong effect of 2HB. These data suggest a shift in red gastrocnemius BCAA homeostasis induced by 2HB treatment.

To compare the effects of 2HB given the quantifiable difference in exercise performance (Fig.2D-G), we adjusted metabolite values based on VO_2_ measures during the exercise protocol or sham control. This adjustment yields values that indicate whether the levels of a given metabolite were high or low relative to the matching VO_2_ measures for each mouse. We observed a number of differences and trends in the red gastrocnemius muscle: reduced levels of BCAA at rest and post-exercise, increased BCKA post-exercise, reduced BCHA post-exercise, and reduced 2HB post-exercise (SFig.2B). In white gastrocnemius we only observe a reduction to BCAA and the BCHA post-exercise (SFig.2C). Each of these effects are weaker in the serum VO_2_-adjusted data (SFig.2D). These data demonstrate that 2HB treatment alters muscle BCAA metabolic balance in excess of the changes to mouse VO_2_ during exercise.

### 2KB is a fuel for oxidative metabolism

The metabolomics analysis led us to hypothesize that physiological 2HB accumulation is associated with competition for the BCAA degradation pathway, limiting oxidation of 2KB. However, despite validation as a substrate for BCKDH in isolated enzyme assays^14^, 2KB has not been validated as a potential fuel for mitochondrial oxidative metabolism. Thus, we next aimed to investigate the potential of 2HB and 2KB as fuel for oxidative metabolism *in vitro* and dependency of 2KB oxidation on the BCAA degradation pathway. We subjected cells to Seahorse assays monitoring the oxygen consumption rate (OCR) during a standard ‘mitochondrial stress test’ protocol following acute administration of 500-5000 μM of either Na2HB, Na2KB, or NaCl control (Fig.3A,B). With C2C12 myoblasts, we found a modest increase to OCR immediately following 2KB addition, with no response to 2HB (Fig.3A,B). Presence of 2HB or 2KB provided a small increase to the maximum OCR. We hypothesized that the standard assay media saturated cells with fuel, leaving little capacity for 2HB or 2KB as a fuel. We therefore repeated the assays with no pyruvate in the medium. Similarly, an acute uptick in OCR was observed upon addition of 2KB, but not 2HB (Fig.3A,B). There was now a dose-dependent increase to maximal OCR in cells treated with 2KB, up to a similar magnitude when pyruvate was included. Addition of 2HB was not able to recapitulate the maximum OCR observed when pyruvate was present in the media. These trends could be replicated with mouse embryonic fibroblasts (MEF), although with a lower magnitude (SFig.3A). These data demonstrate that 2KB is a substantially greater fuel for oxidative metabolism than 2HB.

**Figure 3.**
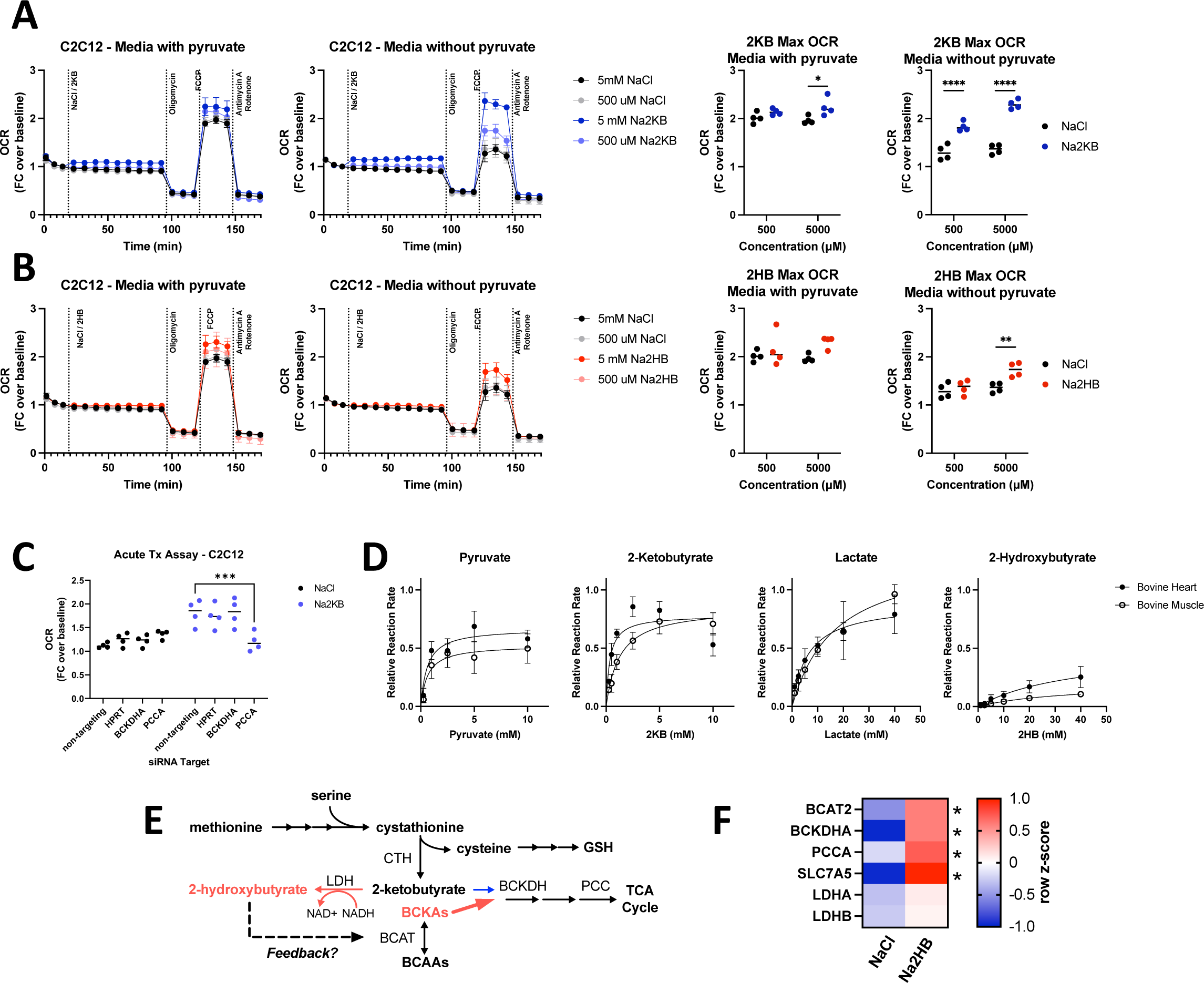
(A,B) Oxygen consumption rate (OCR) during mitochondrial stress test with acute administration of (A) Na2KB or (B) Na2HB as indicated. N = 4 independent assays, mean ± SD. Basal and maximum OCR values displayed with Sidak’s multiple comparisons test displayed, * p < 0.05, **** p < 0.0001. (C) Maximum OCR values from mitochondrial stress tests in C2C12 transfected with siRNA against indicated targets, N = 4 independent assays mean ± SD. Sidak’s multiple comparisons test, *** p < 0.001. (D) Relative reaction rate of LDH assays using indicated substrate. Data are normalized to the maximum reaction rate for a given experiment day for each reaction direction. Data are a summary of five independent experiment days, mean ± SD. (E) Summary figure; we propose that accumulation of 2HB, as occurs post-exertion, is an indicator of saturation in the BCAA degradation pathway, leading to reduction of 2KB instead of mitochondrial oxidation and the potential for metabolic feedback via 2HB. (F) Heatmap summarizing RT-qPCR for C2C12 myoblasts treated with 500 μM of Na2HB or NaCl for 8 hours. N = 4 independent assays, median row z-score displayed with Sidak’s multiple comparisons test, * p < 0.05.

It is possible that the increase to maximal OCR induced by 2KB is due to 2KB promoting glycolysis^20,21^, instead of 2KB being used as a fuel. To investigate the importance of the BCAA degradation pathway for 2KB metabolism, we repeated these experiments in cells transfected with siRNA against *BCKDHA*, *PCCA*, and *HPRT* as an off-target control, or non-targeting siRNA control (SFig.3B). Transfection alone had insignificant effects on maximum OCR, as demonstrated by cells treated with NaCl. Addition of 500 μM 2KB increased maximum OCR as before, however, this was inhibited in cells transfected with siRNA against *PCCA* (Fig.3C). These data suggest that an intact BCAA degradation pathway is required for 2KB promotion of maximal OCR *in vitro*, indicating that 2KB can be used as a fuel for oxidative metabolism.

### 2HB is slowly metabolized by LDH

The poor suitability of 2HB as a fuel relative to 2KB suggests slow conversion to 2KB. Interconversion between 2HB and 2KB is proposed to occur via LDH. Thus, we aimed to quantify the relative rate of 2HB oxidation and 2KB reduction. We conducted activity assays using LDH isolated from bovine skeletal muscle or bovine heart. The former is enriched for LDH isoenzymes with M subunits from *LDHA* expression, while the latter is enriched for LDH isoenzymes with H subunits due to predominant *LDHB* expression. As previously reported, 2KB was an efficient substrate for both LDH enzymes, with a maximum relative reaction rate greater than pyruvate (Fig.3D)^22^. However, 2HB exhibited a maximum reaction rate that was only ∼30% that of lactate using bovine heart LDH, and ∼15% of lactate using bovine skeletal muscle LDH (Fig.3D). These data support a mechanism for the efficient production of 2HB from 2KB, leading to long-lasting 2HB accumulation post-exercise.

Further, we found very low reaction rates for reduction of the BCKA, with a rate between ∼2- 10% the reaction rate of pyruvate. The greatest reaction rate observed was for the keto acid of valine, 2-oxo-3-methylbutyrate, using bovine heart LDH (SFig.3C). No signal was detected above background for the oxidation reaction of any BCHA. These data would be consistent with the relatively low abundance of BCHA, and their slow rate of clearance.

### 2HB metabolic feedback occurs via BCAT inhibition

The slow metabolic processing of 2HB makes it likely to accumulate in cells and trigger a functional response. Based on the effects of 2HB treatment on BCAA, BCKA, and BCHA levels post-exercise (Fig.2), we hypothesized that 2HB provides metabolic feedback to the BCAA degradation pathway (Fig.3E). As an initial test, we found that culturing C2C12 myoblasts with 500 μM of 2HB increased gene expression of multiple factors involved in BCAA degradation, including those involved in 2KB oxidation such as *BCKDHA*, and *PCCA* (Fig.3F).

Based on the structural similarities between 2HB and 2HG, we hypothesized that 2HB may act as a competitive inhibitor of BCAT enzymes. To test this possibility, we conducted activity assays of recombinant human BCAT2 (rhBCAT2) using leucine as a constant substrate and αKG as a varied concentration substrate. In these assays we find a dose dependent reduction to maximal rhBCAT2 activity with increasing concentrations of 2HB in the reaction well (Fig.4A). We conducted similar assays using rhGOT1, an aspartate-oxaloacetate aminotransferase, but found no effect of 2HB on the reaction rate (SFig.4A). Given the modification of BCAT activity, it seemed plausible that 2HB may compete with αKG for active sites in αKGDDs. However, in assays of isolated HIFP4H1 and HIFP4H2, no more than 5% inhibition was observed for concentrations of 2HB up to 10 mM, and similarly little inhibition was observed for assays of isolated KDM6A (STable.1). These data indicate that 2HB inhibits specifically BCAT activity.

**Figure 4.**
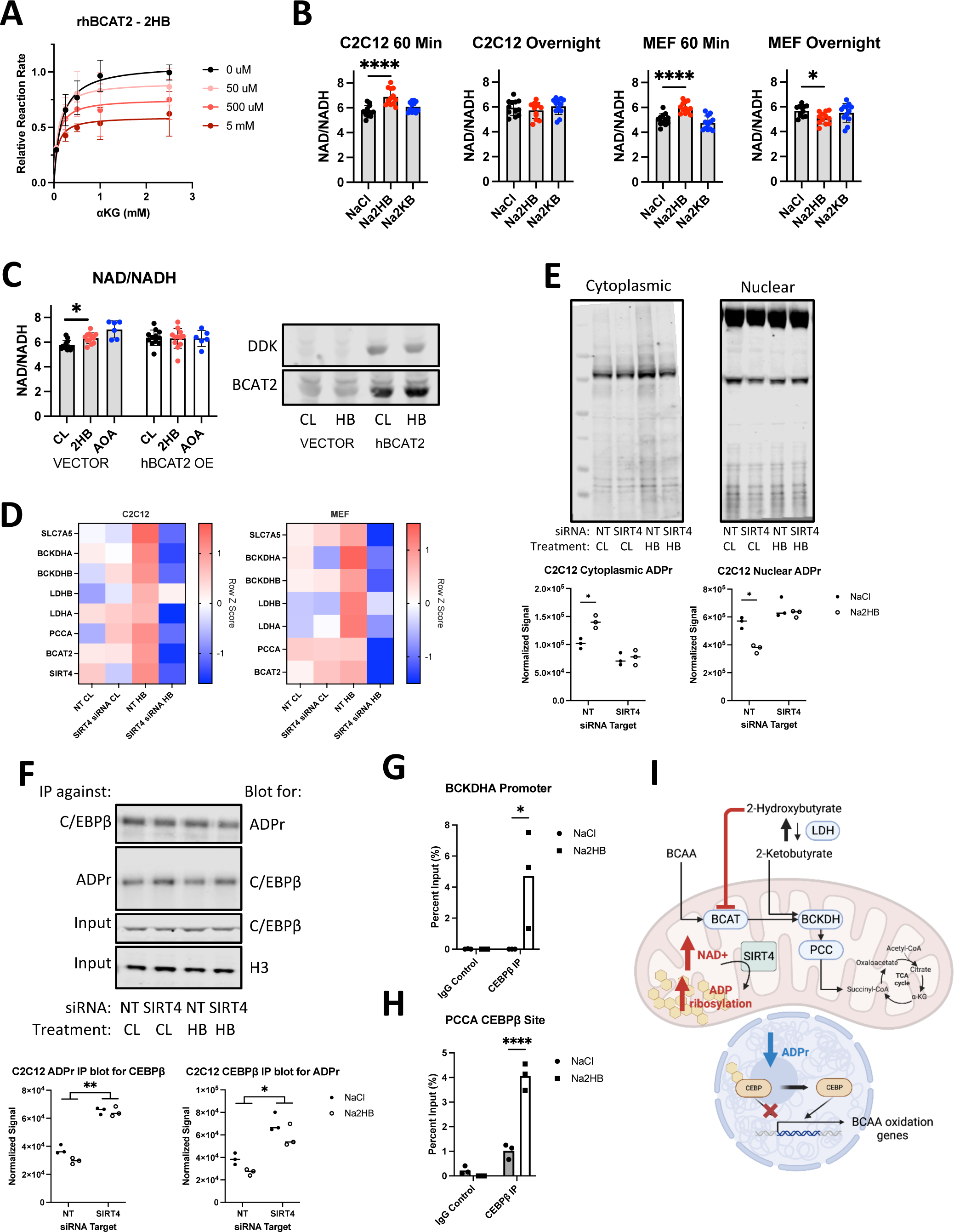
(A) Relative reaction rate for recombinant human BCAT2 activity assays. Leu was kept at a constant concentration of 5 mM, 2HB concentrations are indicated in the legend. Data are normalized to the maximum reaction rate for a given experiment day and data represent three independent experiment days, mean ± SD. (B) Indicated metabolites were added to culture media for indicated time prior to cell harvest for NAD+ and NADH assays, NAD+/NADH ratio displayed. N = 8 independent assays, Dunnett’s multiple comparisons test, **** p < 0.0001, * p < 0.05. (C) C2C12 myoblasts were transfected with vector control or DDK-tagged hBCAT2 before repeating the 60- minute time point NAD+ and NADH assays. 1 mM Aminooxyacetic acid (AOA) included as positive control. N = 12 independent assays, Dunnett’s multiple comparisons test, * p < 0.05. (D) RT-qPCR gene expression for factors related to 2HB and BCAA metabolism in MEFs and C2C12 transfected with siRNA against SIRT4 or non-targeting control, and treated with 500 μM NaCl or Na2HB for 8 hours. N = 4 independent assays, median row z-score displayed. (E) Total protein ADP ribosylation western blots from cytoplasmic and nuclear extracts of C2C12 cells transfected with siRNA against SIRT4 or non-targeting control and treated with 500 μM NaCl or Na2HB for 16 hours. Representative blots shown and quantification from 3 independent experiments. N = 3, Sidak multiple comparisons test, * p <0.05. (F) Immunoprecipitation was conducted on C2C12 nuclear extracts using either antibody against C/EBPβ or poly/mono-ADP ribose, with resultant membranes probed using the opposite antibody. Total C/EBPβ and histone 3 were detected in input samples. N = 3, two-way ANOVA main effect, ** p < 0.01, * p < 0.05. (G) ChIP-qPCR conducted on C2C12 myoblast lysates treated with 500 μM of NaCl or Na2HB for 8 hours. (I) Summary figure displaying proposed mechanism.

Inhibition of BCAT enzymes may induce a range of metabolic disruptions. To broadly investigate cellular response to acute exogenous 2HB, we quantified cellular NAD+/NADH ratio after addition of 500 μM 2HB to culture media. In C2C12 lysates harvested 60 minutes following addition of 2HB we found an increase to the NAD+/NADH ratio, indicating a drop in the cellular reducing potential (Fig.4B). If cells were harvested after overnight culture, NAD+/NADH ratio had rebounded. There was no effect of treating cells with 2KB, indicating that this metabolic response was specific to 2HB. These findings were replicated in MEFs (Fig.4B). To determine if the NAD+/NADH shift induced by 2HB exposure was dependent upon BCAT expression, we transfected C2C12 cells with a vector driving constitutive expression of DDK-tagged hBCAT2, or empty vector control. The increase to NAD+/NADH could be replicated in vector control cells, but the NAD+/NADH ratio in hBCAT2- transfected cells was insensitive to 2HB addition (Fig.4C, SFig.4B). These data describe an acute metabolic shock induced by addition of 2HB to cells, dependent upon BCAT inhibition.

### 2HB treatment stimulates SIRT4 ADP ribosyltransferase activity

Increases to cellular NAD+ are sensed by Sirtuins, a family of deacetylase enzymes that modify cellular metabolism. Sirtuin 4 (SIRT4) is localised to the mitochondria and is previously described to induce increased expression and post-translational activation of the BCAA degradation pathway^23,24^. We hypothesized that SIRT4 would be stimulated by the acute shift in NAD+ induced by exogenous 2HB, and that SIRT4 may drive the cellular response to 2HB. First, we transfected MEFs and C2C12 myoblasts with siRNA against SIRT4 and checked for transcriptional induction of BCAA degradation pathway genes following overnight culture in media supplemented with 500 μM 2HB or NaCl control (SFig.4C). We found induction of genes including *BCAT2*, *BCKDHA*, *BCKDHB*, *PCCA*, *SLC7A5*, as well as *LDHB* and *LDHA* in both MEFs and C2C12s treated with 2HB compared with controls. Transfection with siRNA against SIRT4 disrupted the induction of these genes when cells were treated with 2HB (Fig.4D). These data indicate that expression of SIRT4 is critical for the transcriptional feedback induced by 2HB treatment.

SIRT4 is perhaps better characterized as an ADP ribosyl transferase than as a deacetylase^25^. Addition of ADPr to protein requires consumption of local NAD+. Replacing the consumed NAD+ can cause a compartmental shift in the supply of NAD+, and therefore the source of protein ADPr^26^. Thus, metabolic activity in one compartment may induce transcriptional change via regulating ADPr of transcription factors within the nucleus. To investigate the ability of SIRT4 to regulate compartmental ADPr, we separated cytoplasmic and nuclear extracts of C2C12 myoblasts transfected with siRNA against *SIRT4*, or non-targeting control, treated with 500 μM 2HB or NaCl control for 16 hours. In western blots detecting total poly/mono-ADPr, we see that 2HB treatment increases ADPr in the cytoplasmic compartment, while reducing ADPr in the nuclear extracts, each dependent upon SIRT4 (Fig.4E, SFig.4D). We were next interested in whether 2HB-induced SIRT4 activity might alter the ADPr of a transcription factor relevant to regulation of metabolism. C/EBPβ has been described to regulate adipogenesis^27^, which is also known to involve SIRT4-dependent regulation of BCAA metabolism^24,27^. Importantly, C/EBPβ transcriptional activity was shown to be dependent upon compartmental ADPr, wherein C/EBPβ ADPr reduces its transcriptional activity^27^. Therefore, we conducted immunoprecipitation experiments pulling down total poly/mono-ADPr protein from C2C12 nuclear extracts and blotting for C/EBPβ, or vice versa. We observed a strong effect of SIRT4 knockdown leading to increased ADPr of C/EBPβ, and a trend towards reduced C/EBPβ ADPr in cells treated with 500 μM 2HB for 8 hours (Fig.4F). It is possible that a small shift in ADPr may lead to significant changes to C/EBPβ transcriptional activity. We therefore continued by conducting chromatin immunoprecipitation experiments pulling down C/EBPβ and conducted qPCR for different promoter regions in the genes of *BCKDHA* and *PCCA* containing C/EBPβ consensus binding sites. We found that treating cells with 500 μM 2HB for 8 hours stimulated binding of C/EBPβ to both gene promoters (Fig.4G,H). In summary, we find that treating cells with exogenous 2HB induces a metabolic response via BCAT inhibition, which stimulates SIRT4 to trigger a compartmental change in protein ADPr, leading to C/EBPβ binding to *BCKDHA* and *PCCA* genes (Fig.4I). These data present a mechanism for feedback on the BCAA degradation pathway induced by 2HB accumulation.

### 2HB promotes oxidative metabolism in vitro via the BCAA degradation pathway

We hypothesized that overnight culture with 2HB would increase the expression of BCAA degradation pathway genes and thus promote oxidative metabolism. MEFs and C2C12s were treated overnight with 500 μM of Na2HB, or NaCl control prior to running a mitochondrial stress test or glycolysis stress test to quantify OCR and extracellular acidification rate (ECAR) respectively. Overnight treatment with either 2HB increased both the basal and maximal OCR in MEFs and C2C12s (Fig.5A,B). MEFs exhibited an increased ECAR, indicative of increased glycolytic capacity from treatment with 2HB (SFig.5A). Collectively, these data indicate increased metabolic activity, and particularly an increase to oxidative capacity from overnight culture with 2HB.

**Figure 5.**
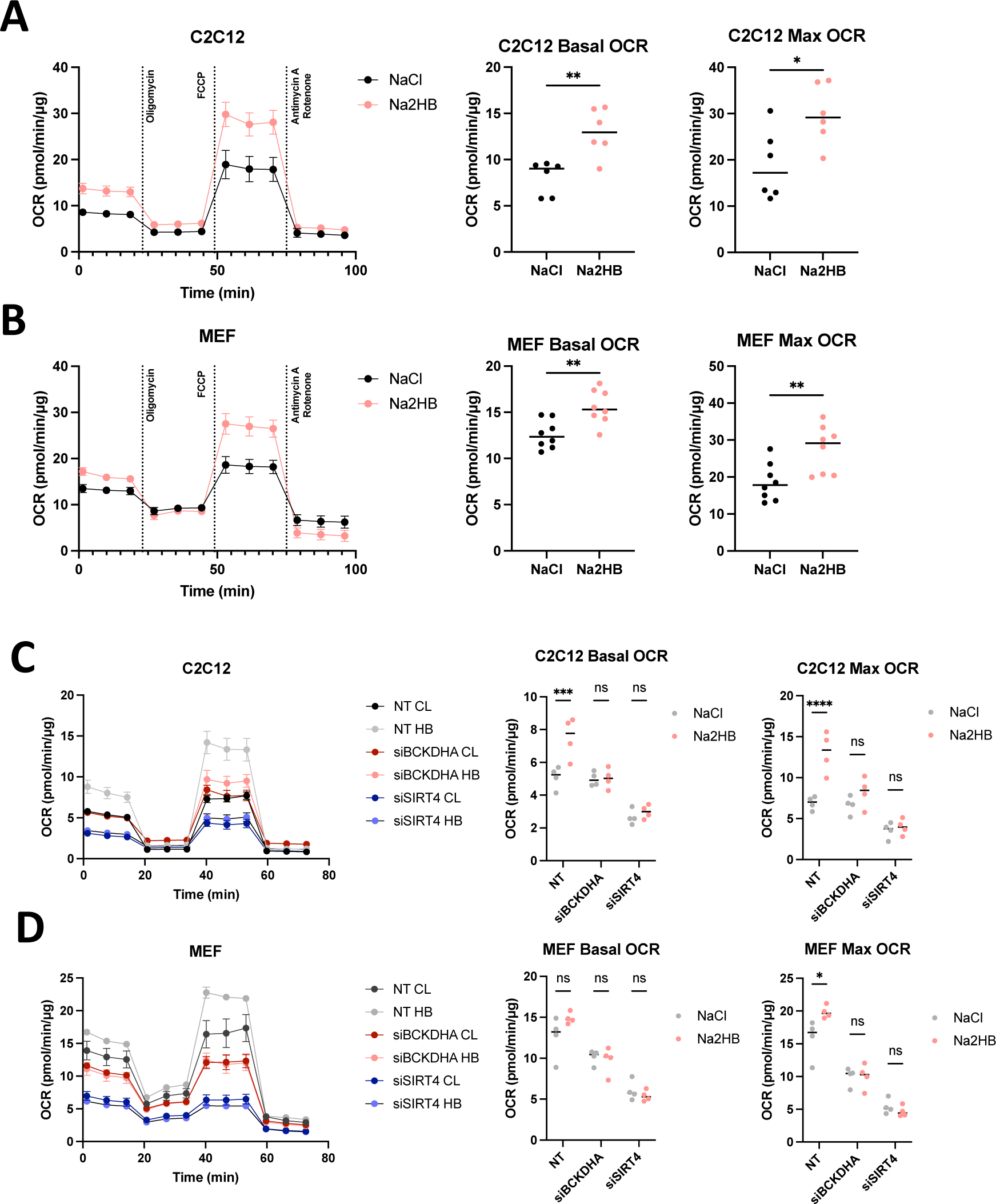
(A) C2C12 and (B) MEFs cultured overnight in media containing 500 μM of Na2HB, or NaCl as indicated, prior to mitochondrial stress test, mean ± SD. Basal and maximum OCR values displayed with Student’s t-test results; N = 6 independent assays ** p < 0.01, * p < 0.05. OCR during mitochondrial stress tests, basal OCR, and maximum OCR for (C) C2C12s and (D) MEFs transfected with siRNA against indicated gene targets. N = 4 independent assays, mean ± SEM Sidak’s multiple comparisons test, **** p < 0.0001, *** p < 0.001, * p < 0.05.

We hypothesized that the increased oxidative capacity induced by 2HB was dependent upon SIRT4 stimulation and activity of the BCAA degradation pathway. To address this possibility, we transfected C2C12s and MEFs with siRNA against SIRT4, BCKDHA, or non-targeting control. Cells were then cultured overnight in media supplemented with 500 μM Na2HB or NaCl, then subjected to a mitochondrial stress test. As before, both MEFs and C2C12s treated with 2HB exhibited an increased basal and maximal OCR (Fig.5C,D). Transfection with siRNA against SIRT4 reduced both basal and maximal OCR and abolished the effect of 2HB (Fig.5C,D). Transfection with siRNA against BCKDHA induced a less drastic reduction to basal and maximal OCR, but also showed no response to 2HB treatment (Fig.5C,D). These data show that SIRT4 and the BCAA degradation pathway, including the early step catalyzed by BCKDHA, is critical to the increased oxidative capacity induced by 2HB treatment.

### 2HB-induced increase to oxidative capacity is maintained with chronic treatment

We were interested in whether the increase to cellular OCR would be maintained with chronic treatment. MEFs and C2C12s were treated with fresh media containing 500 μM of Na2HB or NaCl control daily for three days and assays run on the fourth day. We found chronic treatment with 2HB to result in increased maximum OCR in seahorse assays, to a similar effect size as overnight treatment (SFig.5B). We were interested if other mechanisms could be at play with chronic 2HB treatment. 2HB has been reported to promote cervical tumour cell survival via stimulation of the methyltransferase DOT1L^28^. We therefore investigated the potential for epigenetic regulation as a mechanism to explain the observed change in cellular oxidative capacity. However, we found no change to H3K79me3 signal in isolated histones after three days of 2HB treatment (SFig.5C,D). Neither C2C12s or MEFs exhibited any change to H3K27me3, H3K9me2, or H3K4me3, also suggesting no significant regulation of KDMs from three days of 2HB treatment (SFig.5C,D). Finally, we found no change to the expression levels of respiratory chain protein complexes I-V using an OXPHOS antibody cocktail (SFig.5E,F). These data do not support epigenetic regulation or mitochondrial biogenesis as mechanistic alternatives to SIRT4-dependent upregulation of BCAA degradation pathway for metabolic responses to chronic 2HB treatment.

### Repeated 2HB treatment replicates *in vitro* mechanisms

We next aimed to find evidence for mechanisms leading to the increased oxidative capacity in mice treated with 1 mmol/kg 2HB for one week. We focused analysis on the soleus muscle, as skeletal muscle accounts for the majority of BCAA degradation in the body and the oxidative muscle fibers found in soleus muscle possess greater mitochondrial content for BCAA degradation capacity^29^. We found that soleus muscles from mice treated with 2HB for four days exhibited increased ADPr in cytoplasmic extracts, but reduced ADPr in nuclear extracts (Fig.6A, SFig.6A). We repeated the poly/mono-ADPr and C/EBPβ immunoprecipitation experiments with soleus muscle nuclear extracts. We found that extracts from 2HB-treated mice exhibited reduced C/EBPβ ADPr (Fig.6B). Accordingly, we find the soleus from mice treated with 2HB for one week to exhibit increased gene expression of BCAT2, BCKDHA, and PCCA compared with controls (Fig.6C). We found increased protein expression of PCCA in 2HB-treated soleus lysates (Fig.6D). Further, we found increased BCKDHA protein expression and reduced the amount of BCKDHA phosphorylation at S293 in soleus samples, indicating increased flux through the BCAA degradation pathway (Fig.6E). No changes to BCKDHA expression or phosphorylation were observed in extensor digitorum longus (EDL) lysates (Fig.6F). These data support the mechanisms of 2HB-induced changes to metabolism found *in vitro* as being plausible *in vivo* and support the soleus as a site of response to exogenous 2HB.

**Figure 6.**
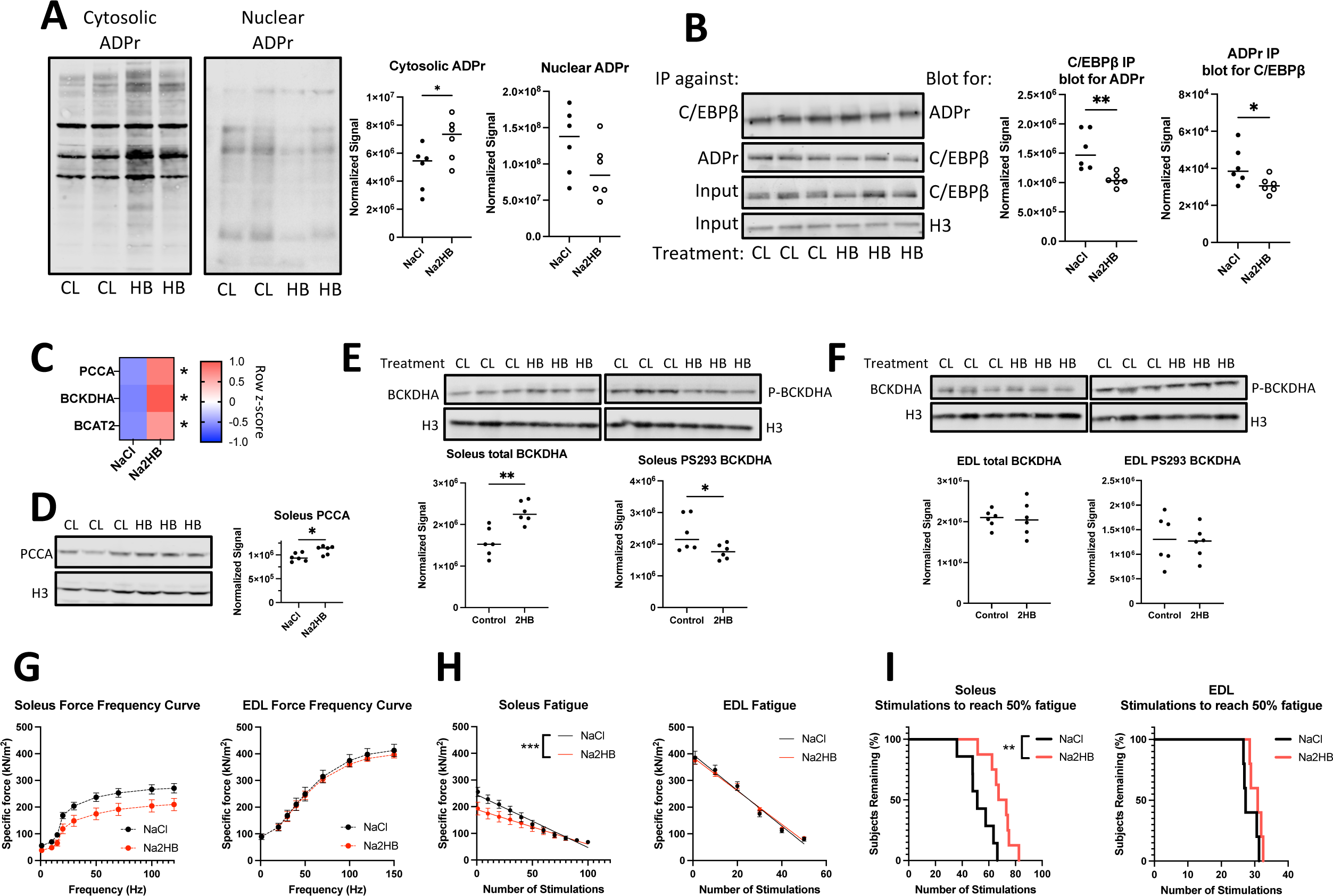
Data come from mice treated for four days with 1 mmol/kg NaCl or Na2HB with harvest of soleus and EDL skeletal muscle on the fifth day. (A) Representative western blots and quantification for total protein ADP ribosylation from cytoplasmic and nuclear extracts of soleus muscle. (B) Immunoprecipitation was conducted on soleus nuclear extracts using either antibody against C/EBPβ or poly-ADP ribose, with resultant membranes probed using the opposite antibody. Total C/EBPβ and histone 3 were detected in input samples. For A,B, N = 6, Student’s t-test, * p < 0.05, ** p < 0.01. (C) RT-qPCR of soleus muscle. N = 4, Sidak multiple comparisons test * p < 0.05. Representative western blots displayed using (D,E) soleus and (F) EDL lysates collected after incremental exercise tests showing total PCCA, BCKDHA, and BCKDHA phosphorylated at S293. N = 10, student’s t-test, * p < 0.05. Ex vivo contractility analysis was conducted on freshly dissected muscle. (G) Force frequency curve. (H) Force production during fatigue protocol with linear regression line displayed. Slopes of linear regression lines are significantly different; F test *** p < 0.001. (I) Time-to-event analysis displaying the number of stimulations in the fatigue protocol for the force output of each muscle to reach 50% of the initial force, log-rank test, ** p < 0.01. N = 7-8 for soleus, N = 5 for EDL. For (G, H) mean ± SEM.

As was observed *in vitro*, there was no change to protein expression of mitochondrial complexes I-V by western blot, indicating mitochondrial biogenesis as an unlikely explanation for the changes to oxidative capacity (SFig.6B). We also assessed histone methylation by western blot and found no change to soleus H3K27me3, H3K79me3, or H3K4me3 after one week of treatment (SFig.6C).

### 2HB improves soleus muscle resistance to fatigue

We hypothesized that the improved performance on exercise tests provided by 2HB was dependent upon changes intrinsic to oxidative skeletal muscle, which exhibits the increased activation and expression of BCAA degradation factors. To investigate, we conducted *ex vivo* contractility analysis of skeletal muscle isolated from mice treated with 1 mmol/kg/day Na2HB or NaCl control for four days. Each muscle was stimulated to produce a force-frequency curve, followed by a fatigue-inducing protocol. Soleus and EDL reached a peak specific force of approximately 200- 270 kN/m^2^ and 400 kN/m^2^ respectively, with a trend towards reduced force production in 2HB- treated soleus (Fig.6G). Soleus muscles from 2HB treated mice demonstrated a slower rate of fatigue; the median number of stimulations required to reduce force production by 50% increased from 51 in controls to 70 in 2HB-treated soleus muscles (Fig.6H,I). EDL from 2HB-treated mice performed similar to controls in the fatigue protocol (Fig.6H,I). These data demonstrate a functional change intrinsic to the skeletal muscle of 2HB-treated mice, specific to oxidative muscle groups.

## Discussion

2HB has to date been described as a waste product that is released into the circulation during periods of metabolic or oxidative stress, for example, in acute infection, or post-exercise^7,9,10^. Here we describe an axis of cellular responses to increased levels of 2HB which regulate the BCAA degradation pathway. We find that exogenous 2HB treatment can recapitulate the effects of short-term exercise training in mice, via improving the fatigue resistance of oxidative skeletal muscle. These data provide insight into novel mechanisms involved in physiological changes to metabolism during exercise training.

The accumulation of 2HB is dependent upon the metabolic balance of 2KB. We demonstrate that 2KB is a suitable fuel for mitochondrial oxidation, dependent upon an intact BCAA degradation pathway (Fig.3). We found that PCC was required for 2KB oxidation, while the BCKDH complex was dispensable. This is in line with data showing that pyruvate dehydrogenase may also oxidize 2KB, although this too has been shown to be dispensable^11,30^. Alternatively, 2KB may be reduced by LDH to 2HB. The relative balance of oxidation versus reduction likely depends on cellular demand for NADH, oxygen availability, and capacity within the BCAA degradation pathway^23–25,31^.

We find that 2HB is a poor substrate for LDH oxidation, supporting the well-described accumulation of 2HB post-exercise, with a slow return to homeostatic levels (Fig. 2,3)^2,7^. Together, these observations are consistent with the accumulation of 2HB as an indicator of saturated capacity for 2KB oxidation via the BCAA degradation pathway. Interestingly, this pattern is mirrored by the BCHA, 2-hydroxyisocaproate, 2-hydroxy-3-methylvalerate, and 2-hydroxyisovalerate in human plasma metabolomics following exercise (Fig.2). The BCHA are rare metabolites, which is consistent with our observation of very low relative reaction rates for the reduction of BCKA by LDH (SFig.3). Further, we were unable to observe significant oxidation of the BCHA in isolated LDH assays (SFig.3), which is consistent with the pattern of accumulation without return to baseline for all the hydroxy acids (Fig.2). Thus, we propose that the reduction of 2KB and BCKA indicate oversaturation of the BCAA degradation pathway, leading to the LDH-dependent production of 2HB or BCHA that are metabolically stubborn and tend to accumulate.

It is reported that the liver is the major producer of circulating 2HB^2^, which would suggest that many tissues receive external 2HB, and that therefore the balance of 2HB metabolism will depend upon 2HB oxidation. LDH isolated from bovine heart had a greater capacity for oxidation of 2HB than LDH isolated from bovine skeletal muscle (Fig.3), suggesting that *LDHB* gene expression may provide an advantage when metabolising exogenous 2HB. Skeletal muscle expression of *LDHB* was previously reported to be induced by exercise training, and to be dependent upon peroxisome proliferator-activated receptor γ coactivator 1α (PGC-1α), and that *LDHB* expression was required for generating an exercise training benefit^32^. Together, this would suggest that improved capacity for metabolising 2HB is common to the progression of endurance training. Since we observed no induction to PGC-1α gene expression or mitochondrial biogenesis (SFig.5,6), treatment with exogenous 2HB does not appear to replicate all exercise training responses in oxidative skeletal muscle. It is possible that 2HB-induced effects on BCAA metabolism occur in an early stage of skeletal muscle endurance training, and that additional signals are required to continue training progression.

We found that the *in vitro* improvement to oxidative capacity induced by 2HB was dependent upon expression of BCKDHA and SIRT4 (Fig.5). SIRT4 has been previously reported to produce greater expression and activation of BCAA degradation pathway members ^23,24^, along with C/EBPβ^27^, making this a logical signaling axis for 2HB response. Increased flux in the BCAA degradation pathway is reflected by a reduced proportion of BCKDHA phosphorylation^31,33^, which we observe in the soleus of 2HB-treated mice post-exercise (Fig.6). BCKDHA phosphorylation by BCKDH kinase is inhibited by BCKA. Thus, the reduced BCKDHA phosphorylation is consistent with the increase to BCKA observed in oxidative red gastrocnemius muscle (Fig.2, SFig.2). That we observe reduced levels of muscular BCHA despite the increase to BCKA levels, without an increase in release of BCKA to the serum, suggests an increase to the oxidative muscle capacity for BCKA oxidation (Fig.2).

Together, these data form a logical feedback loop where accumulation of 2HB provides negative feedback to reduce BCAA degradation at the stage of BCAT. Acutely, this would reduce competition for 2KB in the BCAA degradation pathway by slowing conversion of BCAA to keto acids. A sufficiently large acute dose of 2HB may limit mitochondrial fuel by inhibiting BCAT. This model is consistent with the observed rapid increase to NAD+/NADH ratio in MEFs and C2C12s treated with 2HB in culture (Fig.4). We show that this metabolic effect will lead to transcriptional feedback; an increase BCAA and 2KB oxidation capacity will in the long-term reduce the production of 2HB and BCHA, as observed in red gastrocnemius post-exercise in 2HB-treated mice (Fig.2, SFig.2). That we observe similar transcriptional regulation, changes to BCKDHA phosphorylation, and changes to oxidative metabolism phenotypes *in vivo* suggests that a similar mechanism is occurring in response to intraperitoneal injections of 2HB. In the case of exercise, the physiological accumulation of 2HB would occur during high metabolic demand when mitochondrial fuel supply may be sensitive to the 2HB despite a potentially lower concentration than the exogenous doses used in this study. This is supported by the lack of additive effect on mouse ΔVO_2_max when exogenous 2HB treatment was combined with exercise training (Fig.1).

Overall, this study presents an axis of cellular response to 2HB and presents a functional response to repeated 2HB treatment in the form of improved exercise performance and resistance to fatigue in oxidative murine skeletal muscle. Further work investigating cellular responses to accumulated 2HB are warranted as this metabolite accumulates in the circulation of individuals with infection and metabolic disorders^10,12,34^.

## Materials & methods

### Animals

Male C57/Bl6 mice were purchased from Janvier Labs and housed at Karolinska Institutet under specific pathogen-free conditions. All experiments and protocols were approved by the regional ethics committee of Northern Stockholm. Six-week-old mice were housed for two weeks prior to any experimental procedures to acclimate to the new housing. At experimental endpoint mice were terminated by carbon dioxide. Blood was collected via cardiac puncture and spun in tabletop centrifuge to remove coagulated components, yielding serum. Collected tissue was snap-frozen in liquid nitrogen.

### Incremental exercise protocols

An enclosed chamber treadmill (Columbus Instruments) set at 10° incline was used for all exercise and respiration measurements. Eight-week-old mice were acclimated to the treadmill progressively over four days in a standardized protocol across all mice:

> Day 1: mice are placed on stationary treadmill for 5 min. Starting at 6 m/min for 3 minutes, speed is increased 3 m/min every 3 minutes. Completed after 3 minutes at 12 m/min.

> Day 2: mice are placed on stationary treadmill for 3 min. Starting at 6 m/min for 3 minutes, speed is increased 3 m/min every 3 minutes. Completed after 3 minutes at 15 m/min.

> Day 3: mice are placed on stationary treadmill for 3 min. Starting at 9 m/min for 3 minutes, speed is increased 3 m/min every 3 minutes. Completed after 3 minutes at 18 m/min.

> Day 4: mice are placed on stationary treadmill for 3 min. Starting at 9 m/min for 3 minutes, speed is increased 3 m/min every 3 minutes. Completed after 3 minutes at 21 m/min.

Acclimation was completed at least 48 hours prior to any incremental exercise tests concurrent with measurements of O_2_ and CO_2_ respiration. Incremental exercise tests began with a five-minute settling period, three minutes of measurement with the treadmill at rest, followed by an initial treadmill speed of 6 m/min, after which the speed was increased by 3 m/min every three minutes. Exercise and measurements of oxygen consumption continued until mouse exhaustion, defined as when the mouse refused to run after ten continuous seconds in contact with an electrical grid providing a 1.22 mAmp stimulus at a rate of 2 Hz. Figure 2 exercise held the treadmill at 18 m/min for 30 minutes instead of continuing until exhaustion. Oxygen and carbon dioxide was monitored using a carbon dioxide and paramagnetic oxygen sensor within an OxyMax system (Columbus Instruments). Measurements were collected every 15 seconds. Basal VO_2_ measured while the treadmill was at rest was used to calculate the change to VO_2_ (ΔVO_2_) throughout the exercise protocol. ΔVO_2_max was defined as the maximum increase to VO_2_ above baseline achieved during the incremental exercise protocol.

### Cells

Mouse embryonic fibroblasts (MEFs) were originally derived as described previously^35^. C2C12 myoblasts were a gift from Dr. Jorge Ruas. Cells were cultured in Dulbecco’s modified eagle medium (DMEM) supplemented with 10% fetal bovine serum and penicillin/streptomycin. Cells were cultured with 5% CO_2_, humidity and temperature control, in a cell culture incubator in normoxia (21% O_2_, Sanyo).

### Compounds and drugs

Cell culture media was supplemented as indicated with sodium salts of metabolites, including Na2-hydroxybutyrate and Na2-ketobutyrate and NaCl as a salinity control. Na2HB and NaCl were dissolved in sterile PBS and administered to mice at a concentration of 250 mM with a dosage of 1 mmol/kg. Aminooxyacetic acid (AOA; Sigma) was used as an inhibitor of aminotransferases, and the DOT1L inhibitor EPZ004777 (MedChemExpress).

### Expression plasmids and transfections

MEF and C2C12 cells were transiently transfected using Lipofectamine 2000 (Thermo Fisher) with 20 ng per expression plasmid for 2 x 10^4^ cells per well in a 96-well plate. Cells were treated with transfection mixes for 8-24 hours before further treatment. Knockdown experiments used dicer-substrate siRNA (IDT DNA) against BCKDHA, PCCA, or SIRT4, sequences in Supplementary Information. Knockdown was confirmed by RT-qPCR 24 hours after transfection.

### *In vitro* oxygen consumption rate and extracellular acidification rate

Cellular oxygen consumption rate (OCR) was measured using a Seahorse Extracellular Flux Analyser XF96 (Agilent). MEFs or C2C12s were seeded with 2 x 10^4^ cells per well on the day prior to assay. XF DMEM media (Agilent) was supplemented with 10 mM glucose, 2mM of L-glutamine, and 1 mM Na pyruvate as indicated. Cells were subjected to a “mitochondrial stress test” protocol, involving sequential injection of 2.5 uM oligomycin, 1 uM FCCP, and a combination of 1 uM antimycin A with 100 nM rotenone. Cells were also subjected to a standard “glycolysis stress test” protocol with quantification of the extracellular acidification rate (ECAR), involving sequential injection of 10 mM glucose, 2.5 μM oligomycin, and 50 mM 2-deoxyglucose. Listed concentrations are the final concentrations for each compound. All compounds added in seahorse assays were acquired from Sigma. For experiments acutely treating cells with metabolites within the seahorse instrument, OCR and ECAR values were normalised to baseline measures. For experiments investigating the effects of treatments prior to the assay, OCR and ECAR values were normalized to protein levels within each well quantified by BCA assay (Abcam) conducted after the Seahorse assay.

### Western blotting

Total protein was isolated from cells or mouse tissue using RIPA buffer (Thermo Fisher), or lysates were enriched for histone proteins using Histone Extraction Kit (Abcam), or cytoplasmic and nuclear extracts were isolated using NE-PER Nuclear and Cytoplasmic Extraction Reagents (ThermoFisher), each according to manufacturer instructions. Tissues were homogenized using a glass dounce homogenizer (Active Motif) and kept on ice during homogenization. Proteins were separated in SDS-PAGE, transferred onto PVDF membrane, and probed with antibodies against C/EBPβ (Thermo Fisher, MA1-827), BCKDHA (Thermo Fisher, PA597248), BCAT2 (abcam, ab95976), DDK tag (Origene, TA50011-100), Histone 3 (abcam, ab12079), H3K4me3 (Cell Signaling Technologies, 62255), H3K9me2 (Cell Signaling Technologies, 4658), H3K27me3 (Cell Signaling Technologies, 9733), H3K79me3 (Cell Signaling Technologies, 4260), OXPHOS antibody cocktail (abcam, ab110413), or Poly/Mono ADP-ribose (Cell Signaling Technologies, 83732) and detected using infrared labelled secondary antibody and an ODYSSEY imaging system (LICOR).Antibody signal was normalized to total protein quantified by REVERT total protein stain (LICOR) or appropriate loading control. Western blot data were analyzed using Image Studio Lite software (Image Studio Lite).

### RT-qPCR

Total RNA was extracted using Trizol (Thermo Fisher) according to manufacturer instructions. One microgram of RNA was reverse transcribed using iScript cDNA synthesis kit (BioRad) in a total volume of 20 uL. Real time RT-RT-qPCR was applied to quantify mRNA (7500 Fast Real-Time PCR system, Applied Biosciences Inc., or StepOnePlus Real-time PCR system, Thermo Fisher). Primers are listed in Supplementary Information. Reactions were performed in 96-well MicroAmp Optical plates in duplicate, using SsoAdvanced Universal SYBR Green supermix (BioRad). Gene expression was normalized to *HPRT* unless otherwise indicated.

### Metabolomics analysis

Metabolomics data from Sato et al. and Rundqvist et al. was selected based on relevance to our hypotheses^2,8^. Data was scaled before correlation analysis and partial correlation coefficient calculated to quantify metabolite pair correlations. Correlation analysis was conducted using the MetScape correlation calculator^36^.

Metabolomics related to Figure 2 was performed by Metabolon. Peak area data was subjected to feature-wise normalization using standard scaling or log-transformation to optimize data column normality^37^. Normalization and scaling were conducted separately for serum, red gastrocnemius, and white gastrocnemius data. Exercise and sham treatments for metabolomics experiments occurred inside of sealed treadmills with respiration measurements taken throughout exercise. As treatment altered exercise performance, for some analyses metabolite data was adjusted based on multiple linear regression of respiration parameters. Metabolomics data analysis was conducted using custom python code. Statistical comparisons conducted using statsmodels python package^38^. Heatmaps produced using seaborn python package^39^ or Prism9 (GraphPad).

### LDH, BCAT, KDM6A, Prolyl hydroxylase activity assays

Activity of LDH was assessed using of isolated bovine heart LDH (Sigma), isolated bovine muscle LDH (Sigma), or mouse tissue lysates as described previously^40^. Briefly, one unit of isolated LDH enzyme or 5 μg of tissue lysate was added to a solution of substrate with 2 mM NAD+ for oxidation reactions or substrate with 0.5 mM NADH for reduction reactions, in 20 mM Tris-HCl buffer pH 8.0. Activity of BCAT was assessed using recombinant human BCAT2 (rhBCAT2; R&D Systems)^41^. Briefly, the reaction catalyzed by BCAT2 transferring an amino group from leucine (Leu) to αKG, producing ketoisocaproate (KIC) and glutamate (Glu) respectively, is coupled to the NADH-consuming reaction catalyzed by leucine dehydrogenase from *Bacillus cereus* (LeuDH; Sigma), which converts KIC back to Leu. The reaction mixture consisted of 5 μM pyridoxal phosphate, 50 mM ammonium sulfate, 5 mM DTT, a range of αKG, 10 mM Leu, 0.5 mM NADH, and a range of Na2HB in 100 mM potassium phosphate buffer pH 7.4. One unit of LeuDH along with 100 ng of rhBCAT2 was added to the reaction mixture to initiate the reaction. Upon addition of enzyme, reaction plates were immediately transferred to a plate reader (Synergy Biotek) set at 37°C and absorbance at 340nm quantified every 30 seconds for ten minutes. The initial maximum rate of reaction following any present lag phase was abstracted as the reaction rate.

Activity assays for human KDM6A and the prolyl hydroxylases, HIFP4H1 and HIFP4H2, were conducted as described previously^42–44^.

### Skeletal muscle *ex vivo* contractile function and fatigue

Skeletal muscle was prepared and assessed as described previously^45^. Briefly, the soleus and EDL muscle were excised under dissection microscope from the right hindlimb with proximal and distal tendons kept intact. Excess adipose tissue was manually cleaned from the muscles and tendons were tied with a braided silk thread and mounted in a 15 mL stimulation chamber between a force transducer and an adjustable holder (World Precision Instruments). Muscles were submerged in a Tyrode solution containing (in mmol/L): 121 NaCl, 5 KCl, 1.8 CaCl_2_, 0.1 EDTA, 0.5 MgCl_2_, 0.4 NaH_2_PO_4_, 5.5 glucose, and 24 NaHCO_3_. The force stimulation chamber was set to 31°C and gassed with 95% O_2_ 5% CO_2_ for a pH of 7.4. Each mounted muscle was adjusted to the length at which the highest twitch force was recorded and then allowed to rest for 15 minutes. The force frequency relationship was determined through stimulations at the following frequencies: 1 (twitch), 10, 15, 20, 30, 50, 70, 100, and 120 Hz for soleus, and 1, 20, 30, 40, 50, 70, 100, 120, and 150 Hz for EDL. The stimuli were interspaced by one minute of rest. Upon completion, the muscle was allowed to rest for seven minutes before starting a fatigue protocol consisting of 100 stimulations (70 Hz, 600 msec train duration, 2 sec interval duration) for the soleus or 50 stimulation (100 Hz, 300 msec train duration, 2 sec interval duration) for the EDL. Muscle recovery was determined at 1, 2, 5, and 10 minutes after the final tetanic stimulation at the same stimulation frequency. Absolute force was expressed in millinewton (mN). Specific force (kN/m^2^) was calculated by dividing the absolute force by the muscle cross sectional area, the latter determined by dividing the muscle mass (with tendons removed) by muscle length and density, assuming a density of 1.06 g/cm^3^.

### Statistical Analysis

Data was visualized and statistical analyses conducted in Prism 9 (GraphPad). Statistical tests and replicates are stated in figure legends.

### Funding

This work was funded by the Knut and Alice Wallenberg Scholar Award, the Swedish Research Council (Vetenskapsrådet), and the Swedish Cancer Fund (Cancerfonden). BJW was funded by a Canadian Institutes of Health Research Postdoctoral Fellowship. RSJ was funded by the Principal Research Fellowship from the Wellcome Trust.

## Supplemental Figure Legends

**Supplemental Figure 1.**
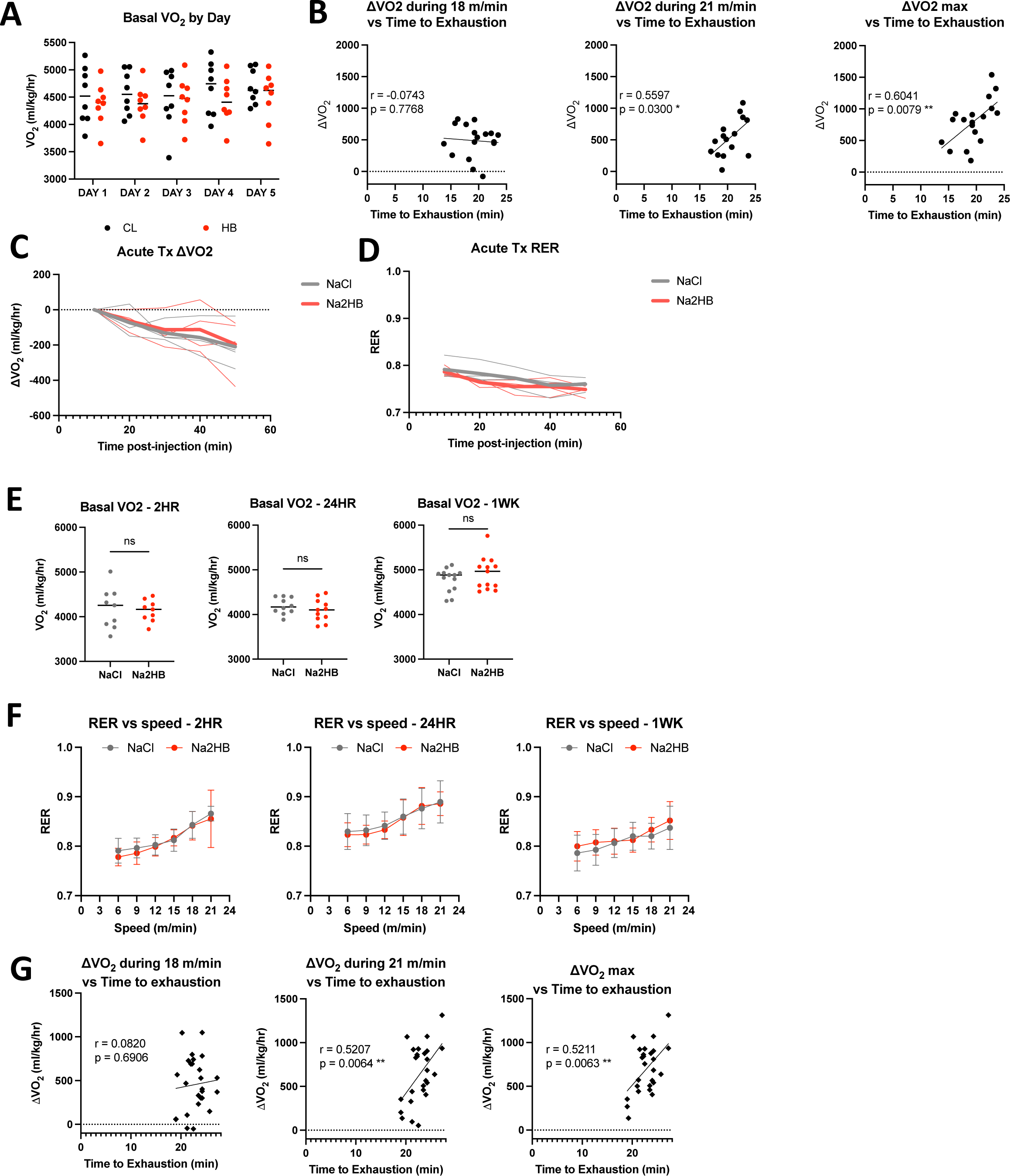
(A) Basal VO_2_ for mice in Figure 1A-E subject to daily incremental exercise tests followed by injection with either 1 mmol/kg of NaCl or Na2HB. Basal VO_2_ is measured on each experiment day during a 5-minute rest period before treadmill begins moving. N = 8. (B) ΔVO_2_ during indicated speed phase or maximum ΔVO_2_ plotted against time-to-exhaustion for mice from Figure 1 A-E on day 5. Pearson correlation coefficients and p values displayed in figure. Data support that mice increase their VO_2_ to match the work rate after the initial boost to VO_2_ above baseline. (C) Mice (N = 3) injected with 1 mmol/kg of NaCl or Na2HB immediately prior to sitting in metabolic chamber at rest show no effect of 2HB on VO_2_ in first 60 minutes. (D) RER for mice in C. For C,D, solid line displays median, each replicate displayed in dashed lines. (E) Basal VO_2_ for mice in Figure 1F-H treated with 1 mmol/kg of NaCl or Na2HB 2 hours, 24 hours, or for four days (label 1WK) prior to exercise test. (F) RER for mice in E, mean ± SEM. For E and F, N = 8-13. (G) Pearson correlation as in B for mice from Figure 1H. N = 13.

**Supplemental Figure 2.**
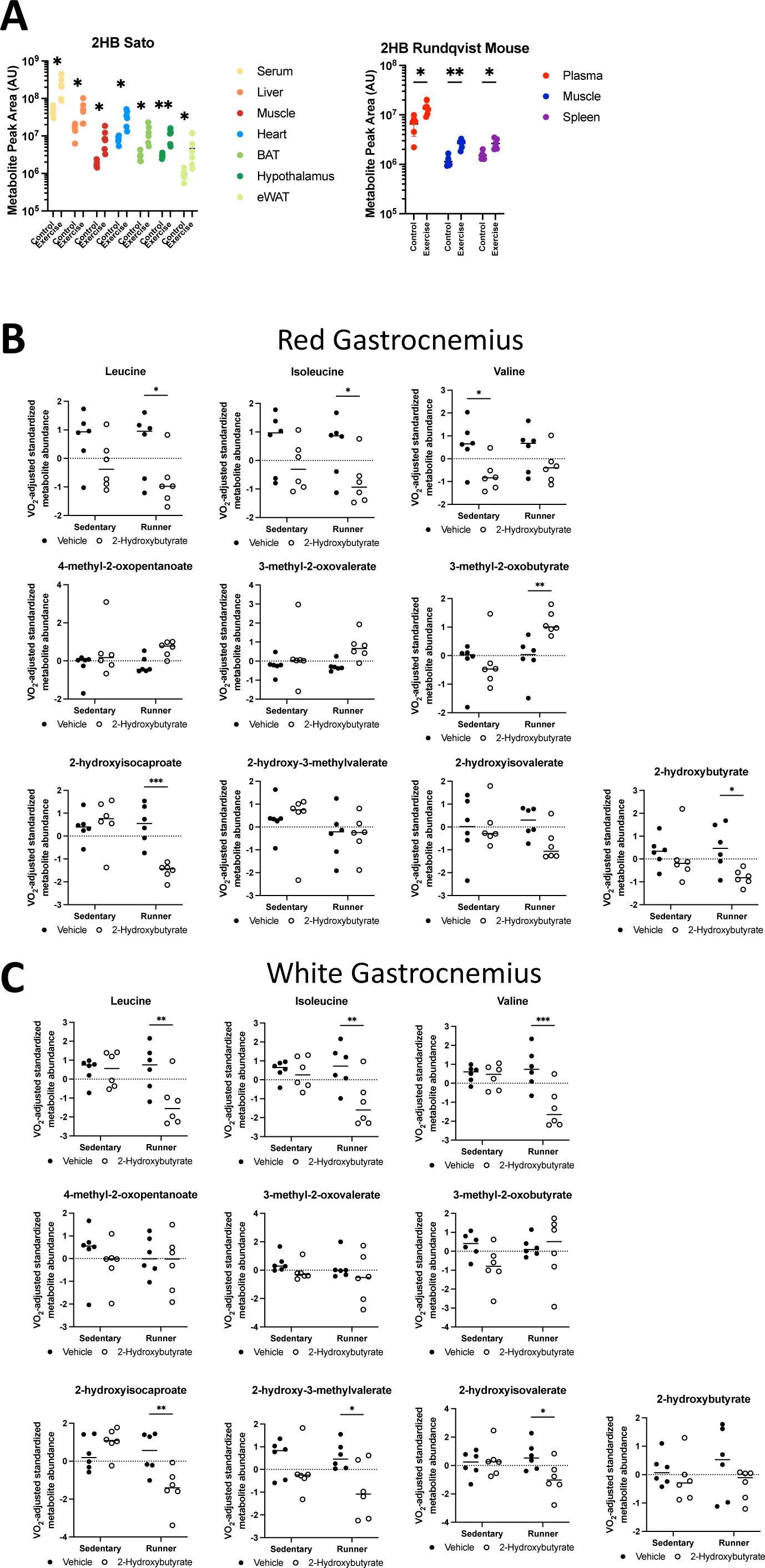
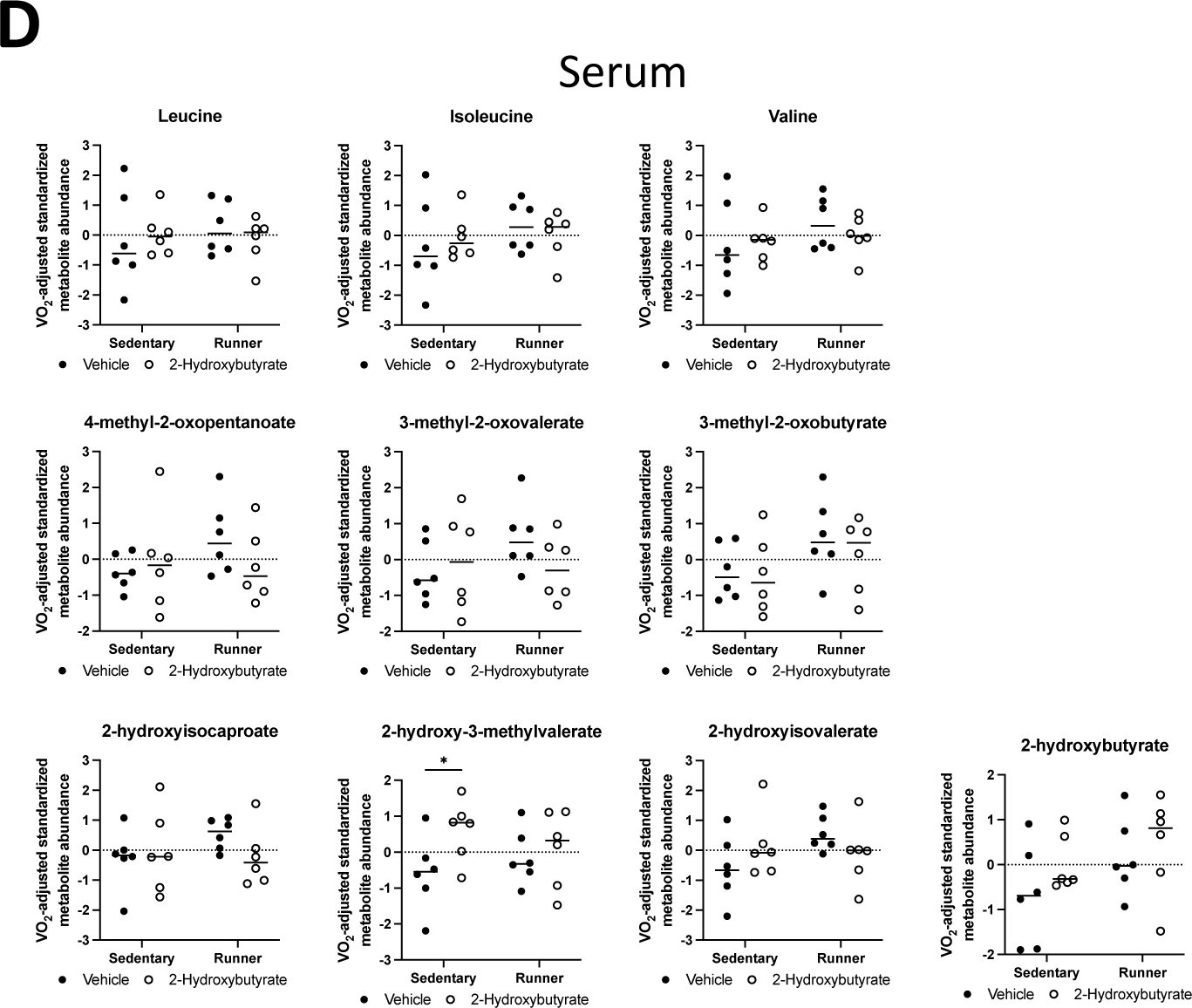
(A) Raw 2HB measures from metabolomics datasets for each organ assessed in Sato et al. and Rundqvist et al. studies. Sidak multiple comparison test results displayed. Plots display VO_2_-adjusted standardized values for the amino, oxo-, and hydroxy- versions of the branched chain amino acids, and 2HB for mice corresponding to Figure 2C-L, for (B) red gastrocnemius, (C) white gastrocnemius, and (D) serum. All data are N = 6. Comparisons made between vehicle and 2HB treated groups of matching exercise condition. Fisher’s LSD test results displayed. * p < 0.05, ** p < 0.01, *** p < 0.001.

**Supplemental Figure 3.**
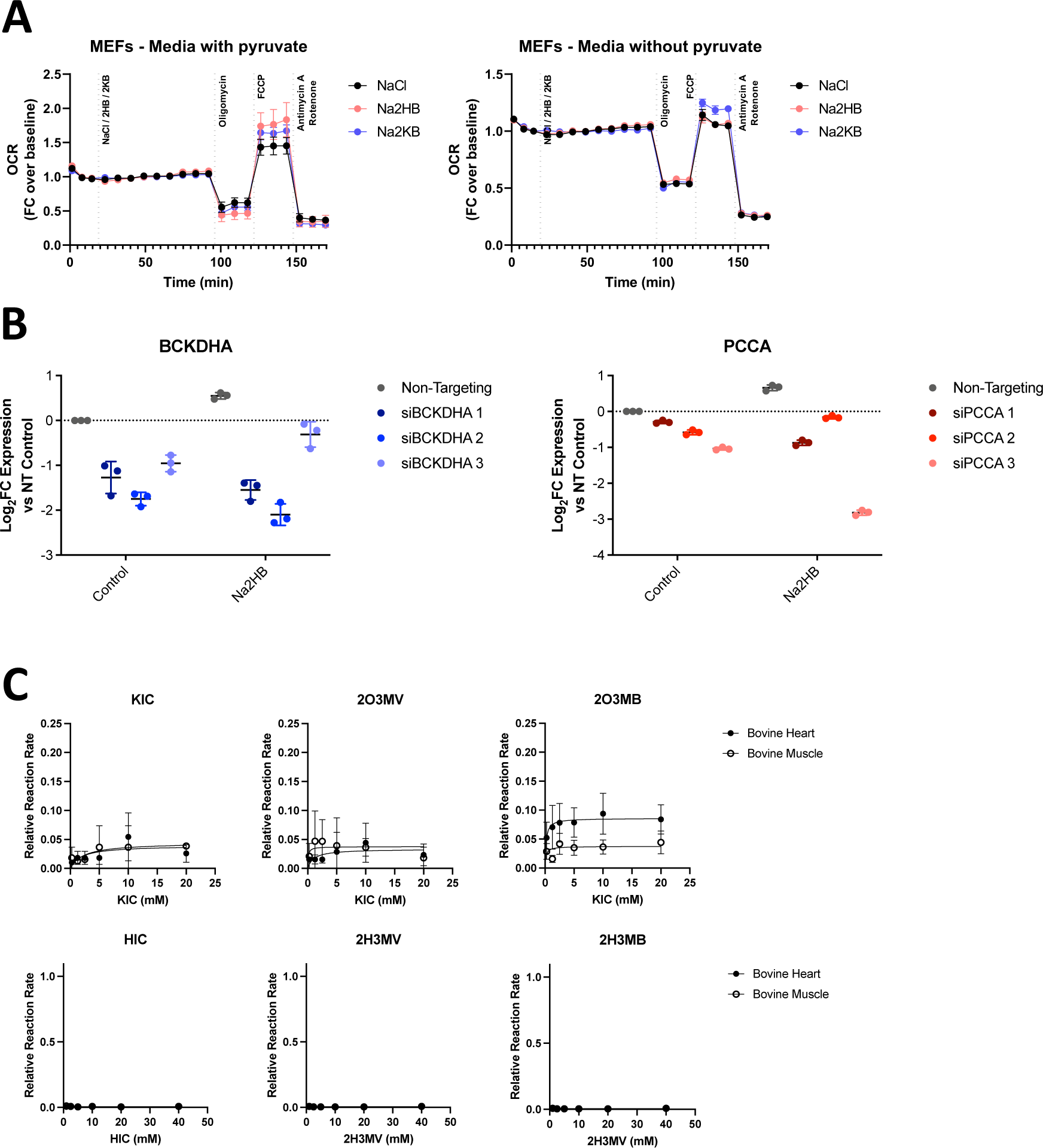
(A) OCR during mitochondrial stress test with acute administration of 500 μM Na2HB or Na2KB as indicated. N = 4 independent assays, mean ± SD. (B) qPCR validation of reduced target gene expression for siRNA transfected cells. Final siRNA selection was based on knockdown of target gene expression in control and 2HB treated cells; siBCKDHA #2 and siPCCA #3. (C) LDH activity assays using keto- and hydroxy-acid versions of BCAA as substrates. From left to right, metabolites are the leucine, isoleucine, and valine derived metabolites; KIC = ketoisocaproate, 2O3MV = 2-oxo-3- methylvalerate, 2O3MB = 2-oxo-3-methylbutyrate, HIC = hydroxyisocaproate, 2H3MV = 2-hydroxy-3- methylvalerate, and 2H3MB = 2-hydroxy-3-methylbutyrate. Data are normalized to the maximum reaction rate in the given reaction direction for the matched experiment day. Data represent mean ± SD of three independent experiment days.

**Supplemental Figure 4.**
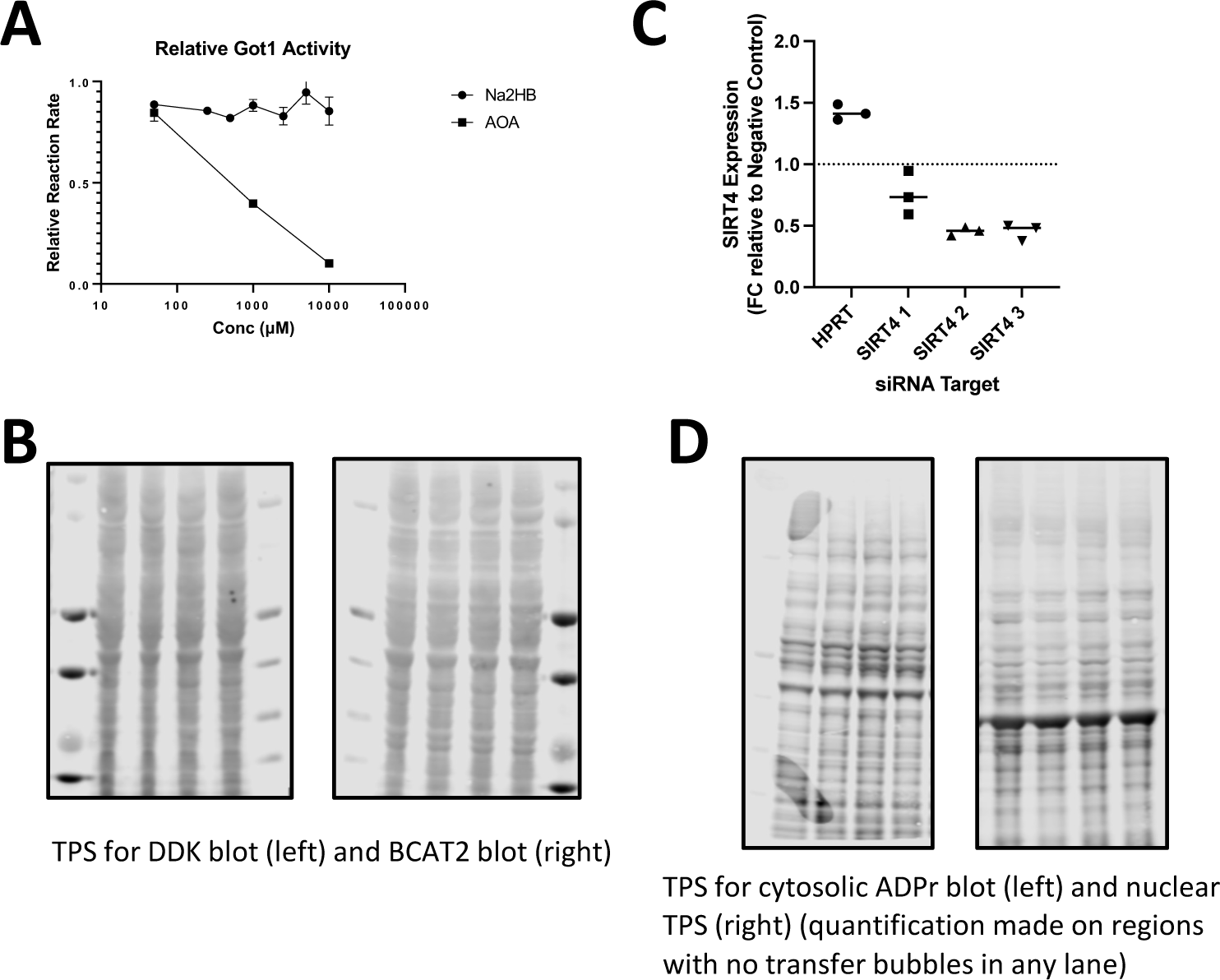
(A) Relative reaction rate for recombinant human Got1 activity. Amino- oxyacetate (AOA) was used as a positive control for Got1 inhibition. Data represent mean ± SD of three independent experiment days. (B) Total protein stains for western blots in Figure 4C. (C) RT- qPCR validation of siRNA against SIRT4. siRNA#3 was selected for experiments in main figures. (D) Total protein stains for western blots in Figure 4E.

**Supplemental Figure 5.**
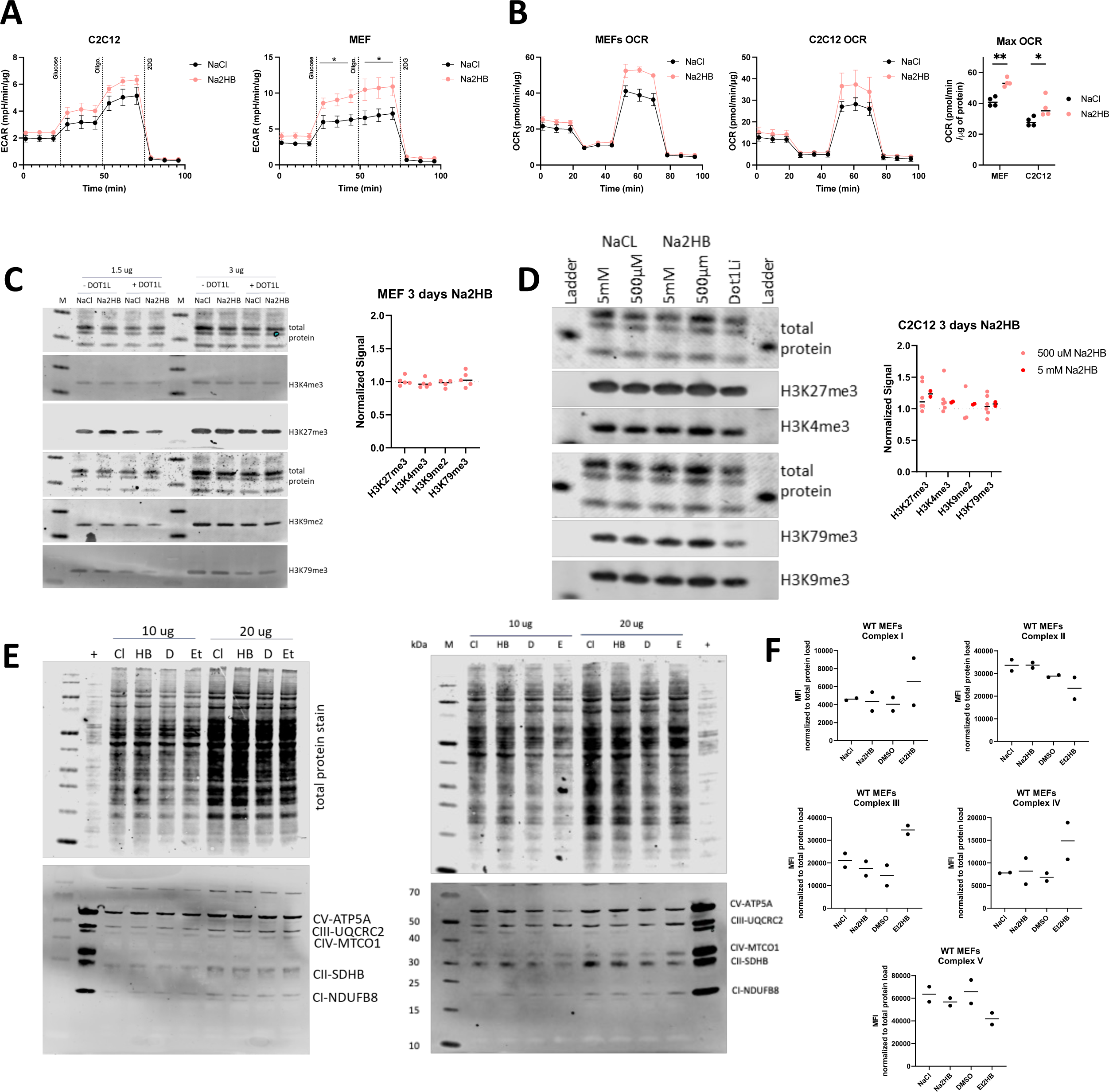
(A) Extracellular acidification rate for C2C12 and MEFs cultured overnight in media containing 500 μM of either NaCl or Na2HB. N = 4 independent assays, mean ± SD. Dunnett’s multiple comparisons test, * p < 0.05. (B) OCR during mitochondrial stress test and maximum OCR for MEFs and C2C12s treated daily with 500 μM of Na2HB or NaCl supplemented media for three days with assays run on day four. N = 4 independent assays, Sidak’s multiple comparisons test, ** p < 0.01, * p < 0.05. (C) Representative western blot and quantification for histones isolated from MEFs treated for three days with 500 μM of Na2HB or NaCl, with or without the DOT1L methyltransferase inhibitor EPZ004777. Amounts of protein loaded are indicated above each blot. (D) Representative western blots and quantification using histones isolated from C2C12 cells treated for 3 days with 2HB or NaCl control at indicated concentrations. EPZ004777 was used as a positive control DOT1L inhibitor. (E) Representative western blot using OXPHOS antibody cocktail from C2C12 (left) and MEF (right) cell lysates following three days of treatment with NaCl (Cl), Na2HB (HB), DMSO (D), or ethyl 2HB (Et). ‘+’ indicates positive control sample. Amount of protein loaded in each well is indicated above. Total protein staining and histone methylation markers displayed. For all western blots, total protein for each well was detected using REVERT total protein stain. (F) Quantification from two independent assays of OXPHOS antibody cocktail western blots using lysates from MEFs treated with 2HB for three days. No observed change to any respiratory chain complex from 2HB treatment is suggested.

**Supplemental Figure 6.**
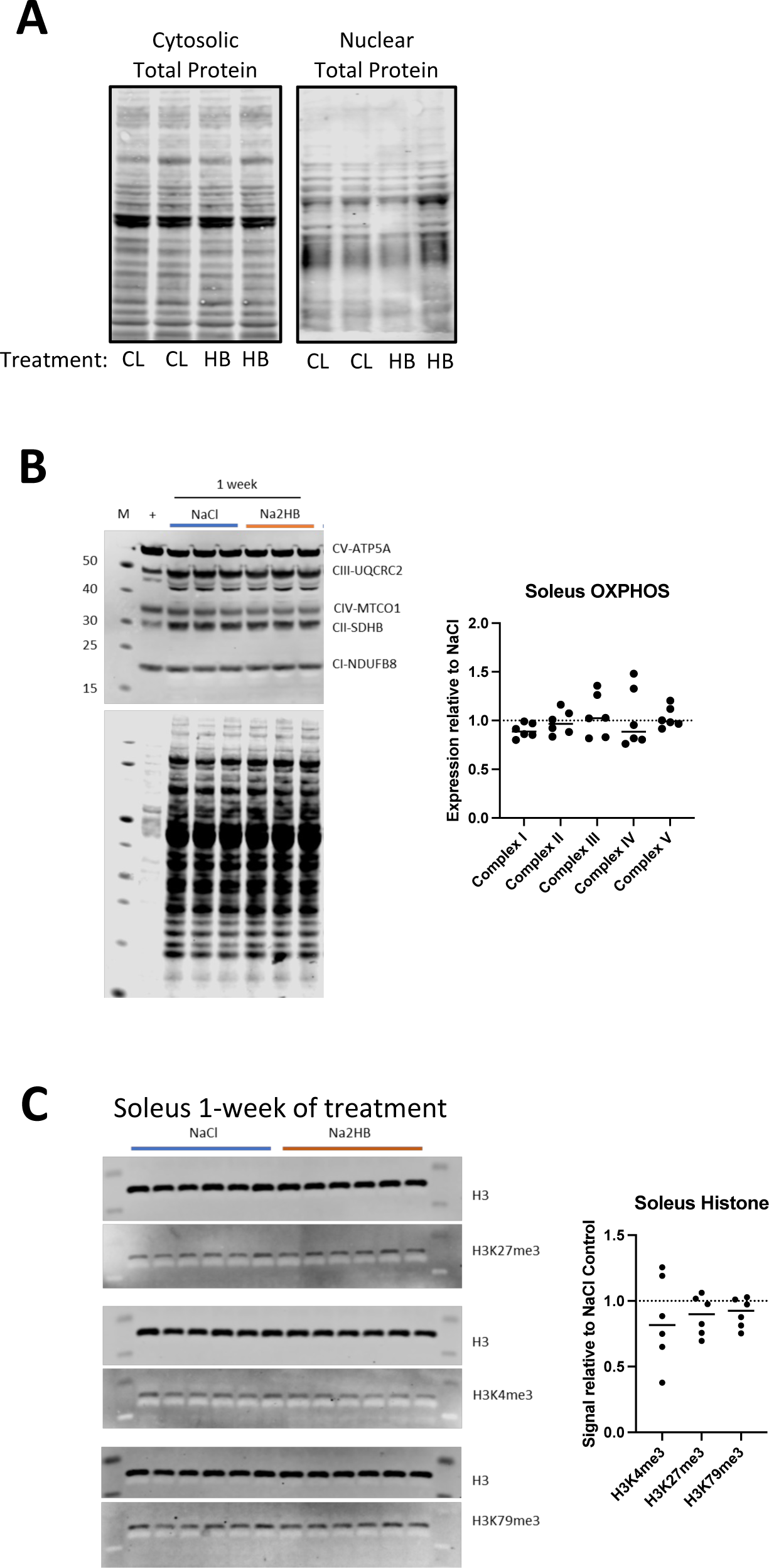
(A) Total protein stain for western blot membrane corresponding to Figure 6A. (B) Quantification and representative western blot using OXPHOS antibody cocktail detection of electron transport chain members in soleus lysates from mice treated with 1 mmol/kg of NaCl or Na2HB for 4 days. (C) Quantification and representative western blot of histone methylation markers in soleus muscle treated as in B. For B and C, N = 6. Data are expressed as fold change relative to average signal for NaCl group on the same membrane.

**Supplementary Table 1.** 2HB and 2KB ICSO for isolated PHO and KDM enzymes

| Enzyme | 2-Hydroxybutyrate |  | 2-Ketobutyrate |  |
| --- | --- | --- | --- | --- |
|  | IC50 (μM) | Inhibition (%) | IC50 (μM) | Inhibition (%) |
| HIFP4H1 | >10000 | <5% | >10000 | <10% |
| HIFP4H2 | >10000 | <5% | >10000 | <5% |
| KDM6A | >10000 | <20% | >10000 | <30% |

*Values are derived from 3 independent assays.*

Wadsworth et al. A 2-Hydroxybutyrate- mediated feedback loop regulates muscular fatigue. Supplementary information. aPCR primer and siRNA sequences.

qPCR primer sequences:

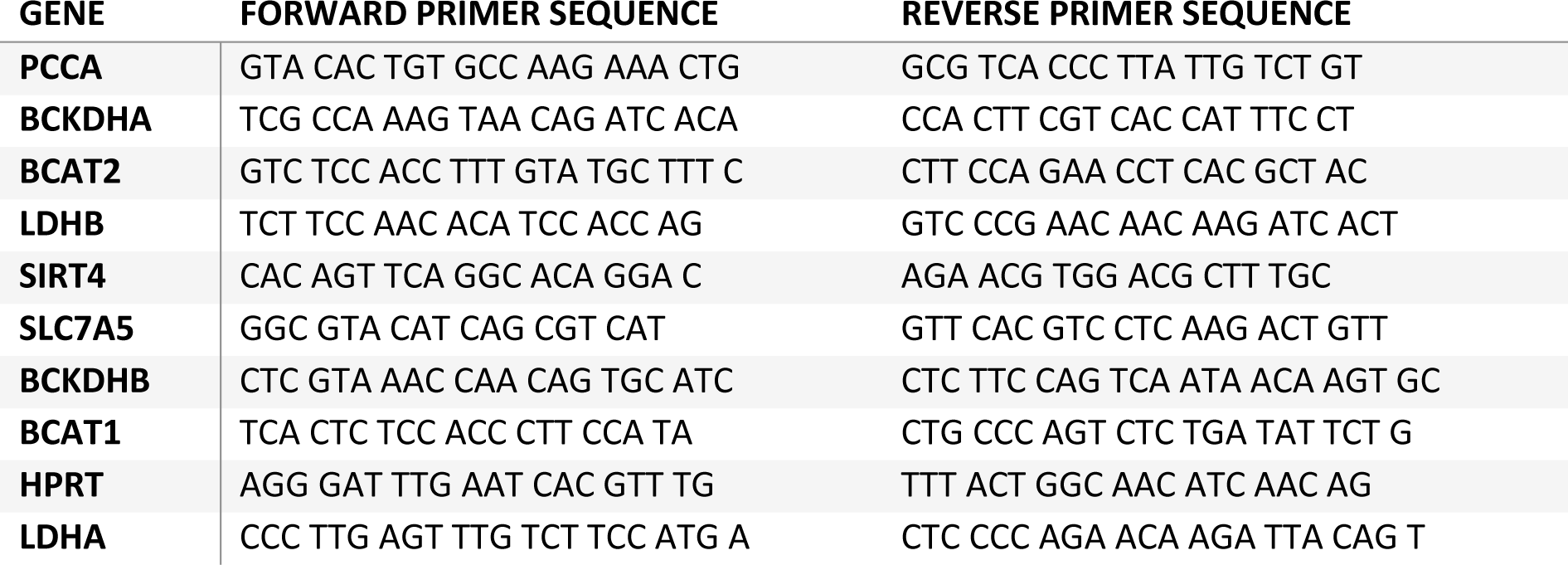

Dicer Substrate siRNA sequences:

| Gene | Sequence 1 | Sequence 2 |
| --- | --- | --- |
| SIRT4 #1 | rArArG rUrCrA rGrUrA rArArC rCrArC rArArC rUrGrU rCrUA T | rArUrA rGrArC rArGrU rUrGrU rGrGrU rUrUrA rCrUrG rArCrU rUrCrU |
| SIRT4 #2 | rCrGrC rUrUrC rArUrU rArGrC rCrUrU rUrCrC rArArG rArAA C | rGrUrU rUrCrU rUrGrG rArArA rGrGrC rUrArA rUrGrA rArGrC rGrCrU |
| SIRT4 #3 | rArArA rArGrC rUrCrC rCrArA rUrArG rCrCrA rUrUrC rUrGA A | rUrUrC rArGrA rArUrG rGrCrU rArUrU rGrGrG rArGrC rUrUrU rUrGrC |
| PCCA #1 | rCrArC rUrUrU rCrArU rUrGrG rArCrC rUrGrA rUrArC rUrCA T | rArUrG rArGrU rArUrC rArGrG rUrCrC rArArU rGrArA rArGrU rGrArC |
| PCCA #2 | rArGrU rArArA rArGrC rUrArC rCrUrC rArArC rArUrG rGrAT G | rCrArU rCrCrA rUrGrU rUrGrA rGrGrU rArGrC rUrUrU rUrArC rUrGrG |
| PCCA #3 | rCrArA rGrCrA rUrUrU rGrUrC rUrArA rArUrA rUrUrA rGrUT T | rArArA rCrUrA rArUrA rUrUrU rArGrA rCrArA rArUrG rCrUrU rGrGrU |
| BCKDHA #1 | rArCrA rCrCrA rArCrC rUrCrU rGrArG rCrArG rUrArC rCrGT G | rCrArC rGrGrU rArCrU rGrCrU rCrArG rArGrG rUrUrG rGrUrG rUrGrG |
| BCKDHA #2 | rArGrA rArGrG rUrCrA rUrGrG rArArG rCrCrU rUrUrG rArGC A | rUrGrC rUrCrA rArArG rGrCrU rUrCrC rArUrG rArCrC rUrUrC rUrUrU |
| BCKDHA #3 | rGrGrG rUrArC rGrGrC rArUrC rArUrG rUrCrA rArUrC rCrGT G | rCrArC rGrGrA rUrUrG rArCrA rUrGrA rUrGrC rCrGrU rArCrC rCrArG |

